# Neuronal LAG3 facilitates pathogenic α-synuclein neuron-to-neuron propagation

**DOI:** 10.1101/2025.01.03.631221

**Authors:** Xiuli Yang, Deok Jeong, Graziella Madeo, Ramhari Kumbhar, Ning Wang, Lili Niu, Junkai Hu, Shuya Li, Kundlik Gadhave, Rong Chen, Fatih Akkentli, Creg J. Workman, Dario A. A. Vignali, Mingyao Ying, Antonello Bonci, Valina L. Dawson, Ted M. Dawson, Xiaobo Mao

## Abstract

Lymphocyte activation gene 3 (LAG3) is a key receptor involved in the propagation of pathological proteins in Parkinson’s disease (PD). This study investigates the role of neuronal LAG3 in mediating the binding, uptake, and propagation of α-synuclein (αSyn) preformed fibrils (PFFs). Using neuronal LAG3 conditional knockout mice and human induced pluripotent stem cells-derived dopaminergic (DA) neurons, we demonstrate that LAG3 expression is critical for pathogenic αSyn propagation. Our results show that the absence of neuronal LAG3 significantly reduces αSyn pathology, alleviates motor dysfunction, and inhibits neurodegeneration *in vivo*. Electrophysiological recordings revealed that αSyn PFFs induce pronounced neuronal hyperactivity in wild-type (WT) neurons, increasing firing rates in cell-attached and whole-cell configurations, and reducing miniature excitatory postsynaptic currents. In contrast, neurons lacking LAG3 resisted these electrophysiological effects. Moreover, treatment with an anti-human LAG3 antibody in human DA neurons inhibited αSyn PFFs binding and uptake, preventing pathology propagation. These findings confirm the essential function of neuronal LAG3 in mediating αSyn propagation and associated disruptions, identifying LAG3 as a potential therapeutic target for PD and related α-synucleinopathies.

## Introduction

Emerging evidence suggests that pathogenic α-synuclein (αSyn) acts as a prion-like seed, inducing the aggregation of endogenous αSyn monomers and facilitating the propagation of pathological features^1^. This process ultimately leads to neurodegeneration and behavioral deficits^2^. Several receptors^3–12^, including lymphatic- activation gene 3 (LAG3), act as key facilitators of the cell-to-cell transmission of pathogenic αSyn both *in vitro* and *in vivo*. Pathogenic αSyn preformed fibrils (PFFs) bind to these receptors and subsequently enter neurons and other cell types via receptor-mediated endocytosis, triggering propagation and neurotoxicity^10–13^. These findings provide critical insights into the mechanisms underlying αSyn PFFs-induced propagation and highlight potential therapeutic targets for developing new treatments for Parkinson’s disease (PD) and related α-synucleinopathies.

Recent research has suggested that LAG3 is not expressed in neurons and does not mediate phosphorylated serine 129 (pS129)-positive αSyn pathology^14^. However, this claim contrasts with other reports indicating that LAG3 is expressed in neurons^10–12, 15–22^ and that LAG3 plays an important role in mediating pathology in α-synucleinopathies^9–12,23^. Public databases show detectable mRNA expression of LAG3 in neurons across multiple species, albert at low levels^15–21^. These expression data that include single cell sequencing analyses, coupled with our recent results^12^ from immunohistochemistry, immunoblot analysis, and RNAscope assays in wild-type (WT) mice versus LAG3 knockout (LAG3^-/-^) mice, as well as the use of a Loxp reporter line with a YFP (yellow fluorescent protein) signal knocked into the LAG3 locus (LAG3^L/L-YFP^)^24^ provide strong evidence that LAG3 is indeed expressed in neurons.

Additionally, the absence of LAG3 significantly reduces pS129-positive αSyn pathology and behavioral deficits induced by the overexpression of human A53T αSyn in transgenic mice^9^. An independent study has shown that LAG3 deficiency in microvascular endothelial cells significantly inhibits αSyn pathology propagation, neurodegeneration, and behavioral deficits^23^. Combined with our previous work on the *in vitro* and *in vivo* role of LAG3 in mediating pathogenic αSyn transmission^10–12^, it is clear that LAG3 plays a crucial role in facilitating the spread of pathogenic αSyn.

Furthermore, we have recently demonstrated that neuronal LAG3 can mediate pathogenic tau neuron-to-neuron transmission using neuronal conditioned knockout mice^22^. However, the *in vivo* role of neuronal LAG3 in mediating α-synucleinopathies remains to be fully tested.

Here, we investigate the role of neuronal LAG3 in mediating the propagation of pS129- positive αSyn pathology, neurodegeneration, behavioral deficits induced by αSyn PFFs inoculation. We assess the basic neuronal activity in the presence of αSyn PFFs in both WT and LAG3^-/-^ mice. Additionally, we evaluate the efficacy of anti-human LAG3 antibody in inhibiting the binding, uptake, and propagation of αSyn PFFs in human dopamine (DA) neurons derived from induced pluripotent stem cells (iPSCs). These studies aim to validate the role of neuronal LAG3 in mediating the propagation of pathogenic αSyn both *in vitro* and *in vivo*.

### Deletion of Neuronal LAG3 Reduces αSyn Pathology Transmission *In Vivo*

To investigate whether neuronal LAG3 is essential for αSyn pathology transmission *in vivo*, we bred LAG3^L/L-YFP^ conditional knockout-reporter mice^24^ (LAG3^L/L-YFP^) with Nestin^Cre^ mice (Jax Lab, strain: 003771)^25, 26^ to generate neuronal LAG3 conditional knockout (LAG3^L/L-N-/-^) mice (Fig. S1A, Fig.1A). Neuronal LAG3 expression depletion was confirmed through this crossbreeding, with immunostaining using anti-LAG3 (LS-B15026) antibody (Fig. S1B, C). Recombinant αSyn PFFs was generated following an established protocol^27^ and confirmed by transmission electron microscopy (TEM) (Fig. S1D). Stereotaxic injection of αSyn PFFs was performed into the dorsal striatum of LAG3^L/L-N-/-^ and LAG3^L/L-YFP^ mice, along with PBS control groups (Fig. 1A).

**Fig. 1.**
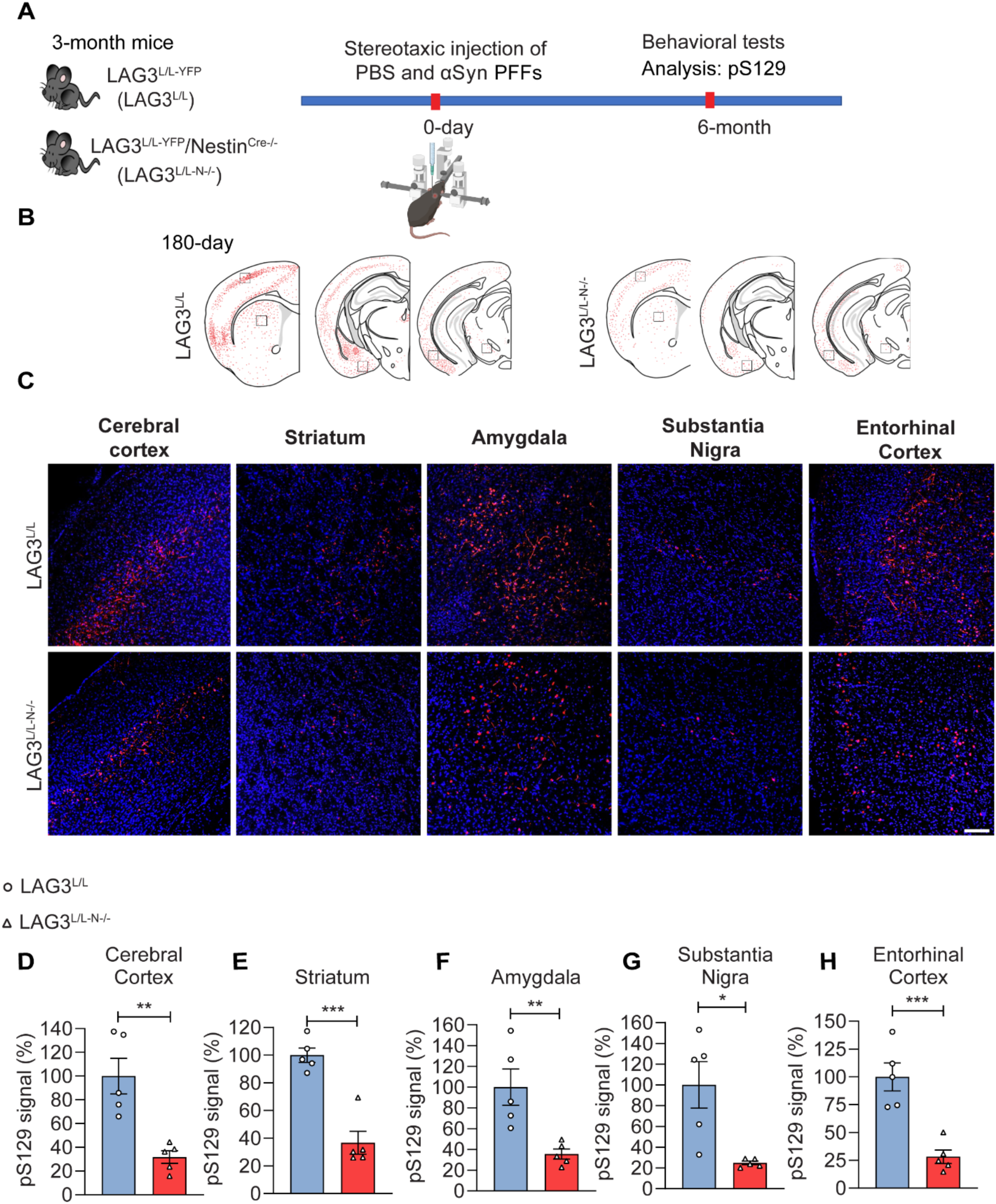
Neuronal LAG3 deletion attenuated αSyn PFFs-induced pathology propagation. **(A)** Experimental design for αSyn PFFs stereotaxic injection and pathology assessment. Immunofluorescence (IF) and immunohistochemistry (IHC) were performed to track pS129-positive αSyn pathology. PFFs were injected into the striatum of LAG3^L/L-YFP^ (control) and LAG3^L/L-N-/-^ (neuronal LAG3 conditional knockout) mice, with comprehensive pathological and behavioral analyses conducted 6 months post-injection. **(B)** Spatial distribution of pS129-positive αSyn pathology following PFFs injection, visualized as red dots across brain sections. **(C)** Quantitative analysis of pS129 immunoreactivity intensity in multiple brain regions, including cerebral cortex **(D)**, striatum **(E)**, amygdala **(F)**, substantia nigra **(G)**, and entorhinal cortex **(H)**. Statistical analysis performed using unpaired two-tailed Student’s *t*-tests. ns, not statistically significant. Data presented as mean ± standard error of the mean (SEM). \**p* < 0.05, \*\**p* < 0.01, \*\*\**p* < 0.001. *n* = 5 biologically independent mice per group. Scale bar, 100 μm.

Two months post-injection, pS129 immunoreactivity was assessed in the entorhinal cortex. Inoculation of αSyn PFFs induced significantly more pS129-positive αSyn pathology in LAG3^L/L-YFP^ mice, whereas LAG3^L/L-N-/-^ mice exhibited a significant reduction of pS129- positive αSyn pathology (Fig. S1 E-H). Six months post-injection, pS129 immunoreactivity was assessed in multiple brain regions, including the cerebral cortex, striatum, amygdala, substantia nigra, and entorhinal cortex (Fig. 1B-H). Robust pS129 signals were observed in these regions of LAG3^L/L-YFP^ mice, whereas pS129 immunoreactivity was significantly reduced in the corresponding regions of LAG3^L/L-N-/-^ mice (Fig. 1B-H). These in vivo findings align with previous observations in LAG3-depleted neurons^10, 12^.

### Deletion of Neuronal LAG3 Alleviates Motor Dysfunction and Inhibits Neurodegeneration Induced by αSyn PFFs

Striatal-injected αSyn PFFs significantly cause motor dysfunction of LAG3^L/L-YFP^ mice in the pole test (Fig. 2A), which is consistent with published studies^10–13^. The behavioral deficits were significantly reduced in the LAG3^L/L-N-/-^ mice (Fig. 2A). The grip strength of forelimbs and four limbs of LAG3^L/L-YFP^ mice injected with αSyn PFFs were significantly reduced (Fig. 2B, C) as published^10–13^. In contrast, deletion of neuronal LAG3 significantly alleviated the reduction of grip strength (Fig. 2B, C). LAG3^L/L-YFP^ mice injected with αSyn PFFs exhibited more latency to fall in the rotarod test (Fig. 2D) similar to prior publication^28^, whereas LAG3^L/L-N-/-^ mice injected mice exhibited no behavioral deficits (Fig. 2D). LAG3^L/L-YFP^ mice inoculated with αSyn PFFs left a flat nest with no shallow walls than PBS-injected LAG3^L/L-YFP^ mice, resulting in a significantly lower nest building scores (Fig. 2E, F) as reported previously^29^. Depletion of neuronal LAG3 significantly enhanced the nest building score (Fig. 2E, F). These results suggest that neuronal LAG3 deficiency significantly alleviates αSyn PFFs-induced motor dysfunction. There was a significant loss of DA neurons as assessed by stereological counting of tyrosine hydroxylase (TH) and Nissl-positive neurons in the substantia nigra of LAG3^L/L-YFP^ mice inoculated with αSyn PFFs (Fig. 2G-I). There was a moderate rescue of DA neurons in αSyn PFFs-injected LAG3^L/L-N-/-^ mice (Fig. 2G-I).

**Fig. 2.**
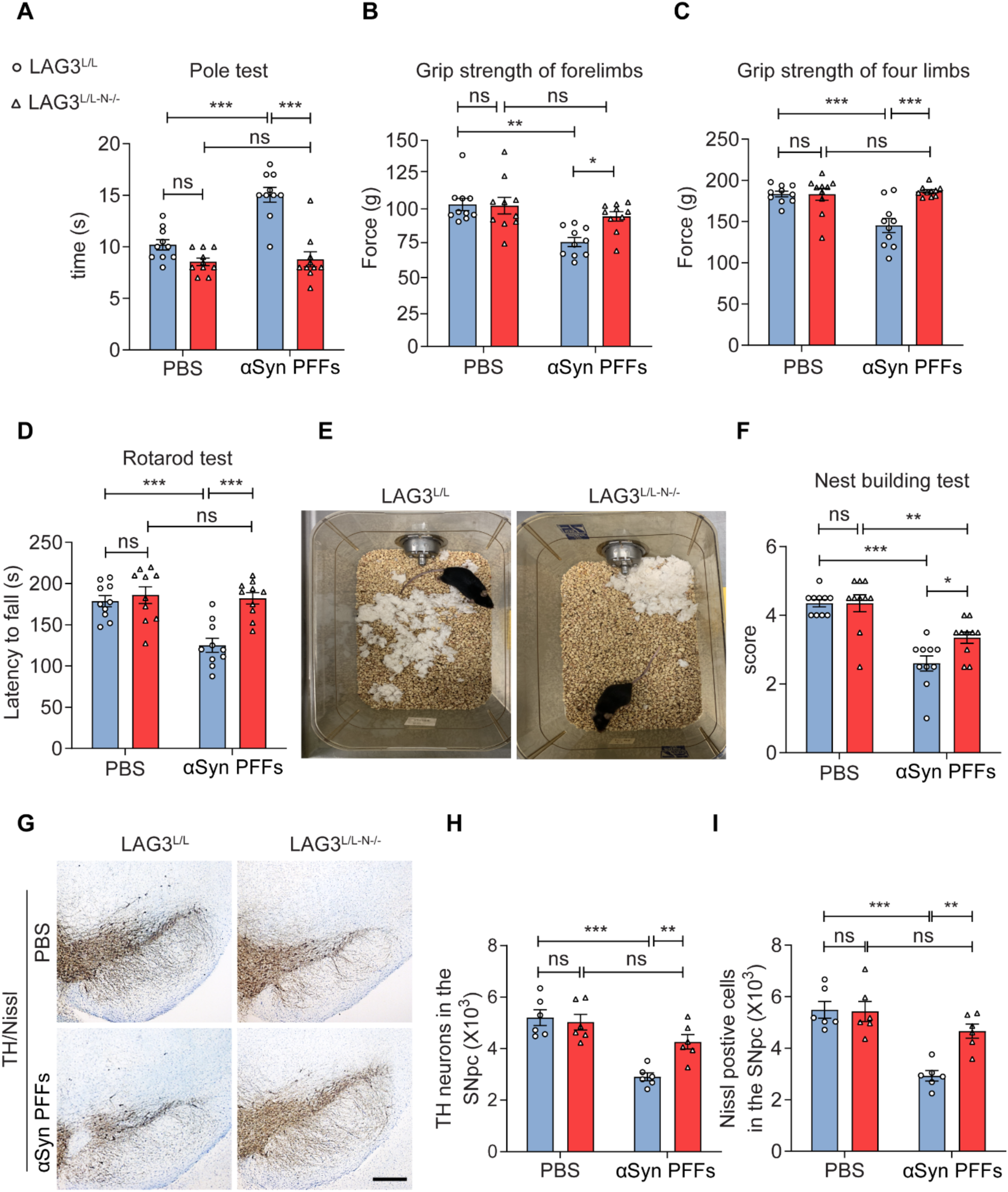
Attenuation of αSyn PFFs-induced behavioral deficits and neurodegeneration in neuronal LAG3 conditional knockout (LAG3^L/L-N-/-^) mice. Behavioral assessments at six months post αSyn PFFs stereotaxic intrastriatal injection: **(A)** Pole test, with a maximum descent time of 60 s to assess motor coordination and bradykinesia. **(B)** Forelimb grip strength test, **(C)** Four-limb grip strength test, **(D)** Rotarod test to evaluate motor performance and balance, and **(E, F)** Representative nest-building images following 16-hour nestlet introduction across experimental groups. Error bars represent mean ± SEM. Statistical significance determined by ANOVA followed by Tukey’s multiple comparisons test. \**p* < 0.05; \*\**p* < 0.01; \*\*\**p* < 0.001; ns, not significant. *n* = 10 biologically independent mice per group. **(G)** Representative photomicrographs of coronal mesencephalon sections showing tyrosine hydroxylase (TH)-positive neurons in the substantia nigra (SN) region. Unbiased stereological quantification of **(H)** TH-positive and **(I)** Nissl-positive neurons in the SNpc region. Data presented as mean ± SEM; \*\**p* < 0.01, \*\*\**p* < 0.001, ns, not significant, *n* = 6 biologically independent mice. Scale bar, 500 μm.

### LAG3 Expression Increases During Aging

Proteome studies have shown that soluble LAG3 levels increase with aging and in an Alzheimer’s disease (AD) mouse model^30^. Brain lysates from young (3 months) and aged (16 months) WT mice were analyzed by immunoblotting with anti-LAG3 antibody (LS-B15026). The results revealed significantly higher levels of LAG3 expression in the brains of aged mice (Fig. S2A, B, C). This aging-induced increase in LAG3 expression was further confirmed through immunostaining, which showed colocalization of LAG3 with NeuN-positive (neuronal nuclei) (ab134014) neurons (Fig. S2D, E).

### Neuronal Intrinsic Excitability and Synaptic Properties in Response to αSyn PFFs

To further investigate the role of neuronal LAG3 in the presence of αSyn PFFs, we performed electrophysiological recordings on acute horizontal midbrain slices from adult LAG3^-/-^ and WT mice. These recordings specifically examined the intrinsic excitability properties of DA neurons within the substantia nigra pars compacta (SNc). Following a 90 minutes *in vitro* incubation with PBS and αSyn PFFs, as previously described^31–34^, SNc DA neurons were successfully filled with neurobiotin during recordings. Post hoc analysis confirmed their dopaminergic identity based on TH immunoreactivity (Fig. 3A, B). Neurobiotin labeling further enabled precise anatomical mapping of DA neurons within the midbrain. Dopaminergic neurons from SNc presented no alterations of intrinsic membrane properties in both WT and LAG3^-/-^ mice when exposed to αSyn PFFs or control condition, where vehicle PBS was used. Exposure to αSyn PFFs significantly increased the firing rate of SNc DA neurons compared to PBS conditions.

**Fig. 3.**
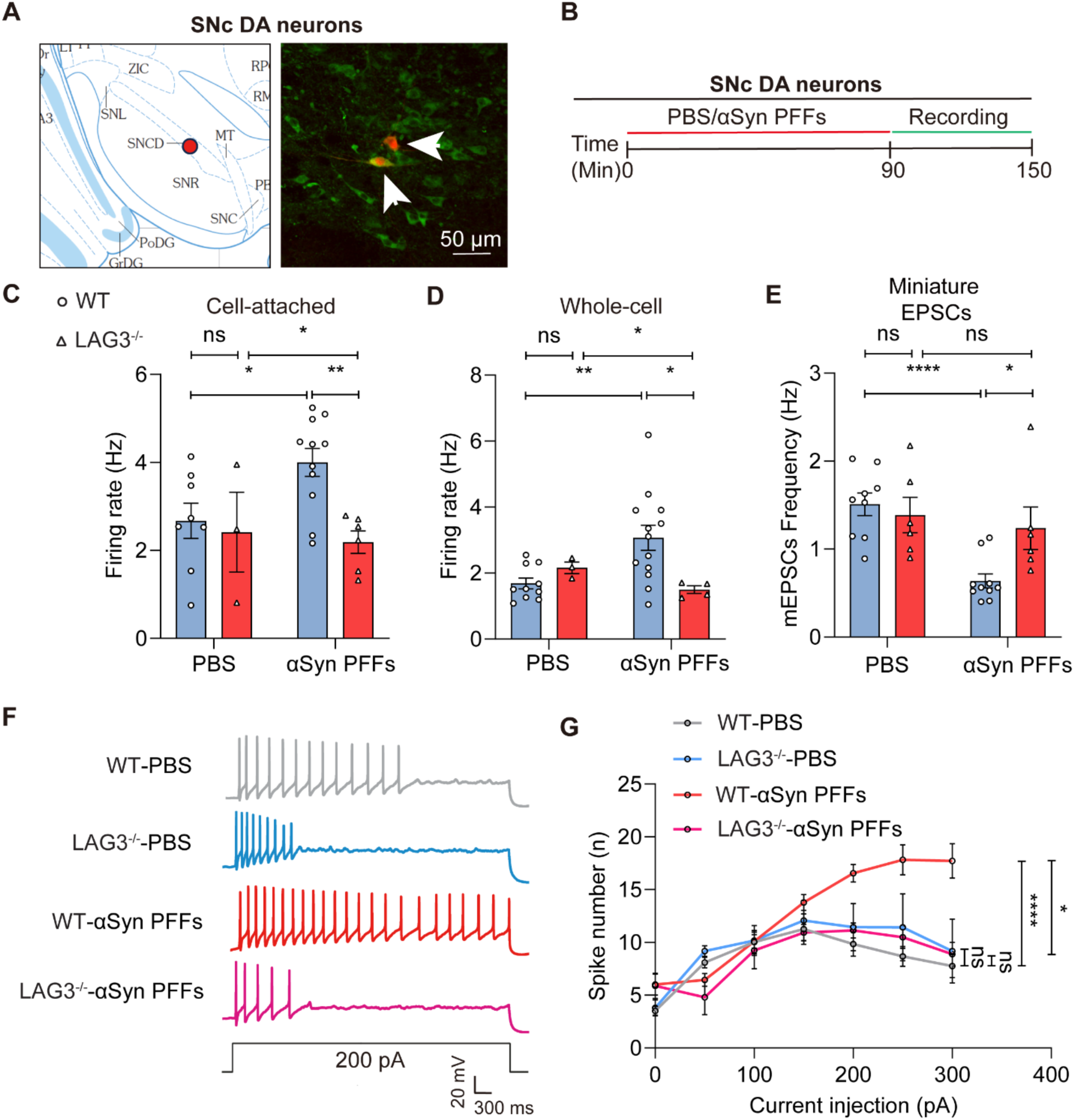
αSyn PFFs increased firing frequencies in SNc DA neurons from WT mice, but not in LAG3^-/-^ mice. **(A-B)** Schematic representation of midbrain slice incubation protocol. **(A)** Left, DA neuron location in SNc. Right: As the white arrow points, neurobiotin-filled cell (red) expressing tyrosine hydroxylase (TH, green) from SNc. Scale bar: 50 μm. **(B)** Experimental timeline for phosphate-buffered saline (PBS) or αSyn PFFs incubation in SNc slices and subsequent electrophysiological recordings. **(C)** Firing frequency of SNc DA neurons in PBS and αSyn PFFs-treated slices from WT and LAG3^-/-^ mice using cell-attached recording. WT mice-PBS, *n* = 8; LAG3^-/-^-PBS mice, *n* = 3; WT mice-αSyn PFFs, *n* = 11; LAG3^-/-^ mice-αSyn PFFs, *n* = 6. **(D)** Whole-cell recording of SNc DA neuron firing frequency in PBS and αSyn PFFs-treated slices from WT and LAG3^-/-^ mice. Note the significant increase in firing frequency in the αSyn PFFs-treated WT group compared to PBS, whereas LAG3^-/-^ mice do not exhibit this increase. WT mice-PBS, *n* = 9; LAG3^-/-^-PBS mice, *n* = 3; WT mice-αSyn PFFs, *n* = 13; LAG3^-/-^ mice-αSyn PFFs, *n* = 4. **(E)** Miniature EPSC frequency in SNc DA neurons from PBS and αSyn PFFs-treated WT and LAG3^-/-^ mice using whole-cell recording. WT mice-PBS, *n* = 9; LAG3^-/-^-PBS mice, *n* = 6; WT mice-αSyn PFFs, *n* = 10; LAG3^-/-^ mice-αSyn PFFs, *n* = 6. **(F)** Representative traces of spontaneous firing activity and neuronal response to hyperpolarizing 200 pA current injection (2 s duration) following PBS and αSyn PFFs *in vitro* incubation. Scale bars: 20 mV and 200 ms. **(G)** αSyn PFFs induces increased SNc DA neuron excitability in WT mice, demonstrated by reduced current accommodation during incremental current injections (50, 100, 150, 200, 250, 300 pA). LAG3^-/-^ SNc DA neurons treated with PBS or αSyn PFFs maintain current accommodation similar to WT control conditions. WT mice-PBS, *n* = 15; LAG3^-/-^-PBS mice, *n* = 4; WT mice-αSyn PFFs, *n* = 11; LAG3^-/-^ mice-αSyn PFFs, *n* = 4. Values expressed as mean ± SEM. \**p* < 0.05, \*\**p* < 0.01, \*\*\*\**p* < 0.0001, ns, not significant. Statistical significance determined by ANOVA with Tukey’s post-hoc correction.

This firing rate enhancement was observed consistently in both cell-attached and whole-cell recording configurations (Fig. 3C, D).

To complement our investigation of intrinsic excitability, we next examined the synaptic transmission properties of SNc DA neurons. Synaptic inputs, particularly excitatory postsynaptic currents, play a critical role in shaping neuronal firing patterns and responsiveness. We assessed both spontaneous (sEPSC) and miniature (mEPSC) excitatory postsynaptic currents to determine how αSyn PFFs exposure affects presynaptic and postsynaptic components of excitatory synaptic activity.

WT neurons exposed to αSyn PFFs exhibited a reduction trend in sEPSC frequency compared to control neurons treated with PBS. Conversely, LAG3^-/-^ neurons did not show any significant change in sEPSC frequency under αSyn PFFs conditions (Fig. S3I). Notably, sEPSC amplitude remained unchanged across all groups, suggesting that the observed frequency alterations are due to presynaptic mechanisms rather than changes in postsynaptic receptor function. (Fig. S3J). mEPSC frequency was significantly reduced in WT neurons treated with αSyn PFFs compared to controls (Fig. 3E). In contrast, LAG3^-/-^ neurons showed no significant differences in mEPSC frequency between control and αSyn PFFs conditions. mEPSC amplitude also showed low variability across all groups, further supporting a presynaptic origin of the frequency changes (Fig. S3K).

We further investigated the intrinsic excitability of SNc DA neurons by measuring the number of action potentials generated in response to a series of current injections. SNc DA neurons exposed to αSyn PFFs showed a significantly higher spike count compared to neurons in control conditions, while the intrinsic membrane properties and intrinsic excitability properties show no significant changes across all groups (Fig. S3A—H).

Moreover, αSyn PFFs-treated neurons displayed reduced accommodation, evidenced by sustained firing at higher current injections. Notably, the spike number was higher across all current steps from 0 pA to 300 pA in αSyn PFFs-treated neurons compared to control. Conversely, SNc DA neurons from LAG3^-/-^ slices exhibited the accommodation phenomenon even when exposed to αSyn PFFs, maintaining a spike pattern similar to control neurons (Fig. 3F, G). These results suggest that LAG3 plays a critical role in the αSyn PFFs-induced excitability changes in SNc DA neurons. The intrinsic membrane properties of SNc DA neurons were not affected by the αSyn PFFs exposure and they were comparable in all experimental groups.

These results indicate that αSyn PFFs enhances presynaptic excitatory drive in WT SNc DA neurons, contributing to increased synaptic input. The absence of these effects in LAG3^-/-^ neurons suggests that LAG3 is a critical mediator of αSyn PFFs-induced synaptic alterations.

### Anti-Human LAG3 Antibody Inhibits the Binding of αSyn PFFs to Human DA Neurons

Given the evidence that mouse neuronal LAG3 facilitates αSyn PFFs-induced propagation *in vivo*, we subsequently investigated the role of human neuronal LAG3 in mediating interactions with αSyn PFFs. Human DA neurons were differentiated from induced pluripotent stem cells (iPSCs) following the protocol outlined in our previous publication^35^. The purity of DA neurons was confirmed to be over 90%, as determined by immunostaining for tyrosine hydroxylase (TH) and NeuN double-positive cells, consistent with the results reported in our earlier work^35^ (Fig. S4A, B).

To investigate the interaction between αSyn PFFs and neuronal binding sites, we employed a cell surface binding assay^10–12, 22^. This assay is widely used to evaluate the interaction between cellular receptor and ligand^21, 36, 37^. Anti-human LAG3 antibody (17B4) and the negative control mouse immunoglobulin G (mIgG) were treated into the human iPSCs-derived DA neuron cultures for two hours incubation. Human iPSC-derived DA neurons were treated with biotinylated αSyn PFFs at varying concentrations (Fig. 4A, B). By following the established protocol of cell surface binding assay^10–12, 22^, binding was detected using streptavidin-alkaline phosphatase staining. Binding analysis revealed that biotinylated αSyn PFFs bound to human DA neurons treated with mIgG with a dissociation constant (KD) of 962.4 nM (Fig. 4A, B). In contrast, treatment with 17B4 significantly reduced binding intensity (Fig. 4A, B). Specific binding of αSyn-biotin PFFs to LAG3 was calculated by subtracting the binding observed in 17B4-treated neurons from that in mIgG- treated neurons, revealing a KD of 820.8 nM (Fig. 4B).

**Fig. 4.**
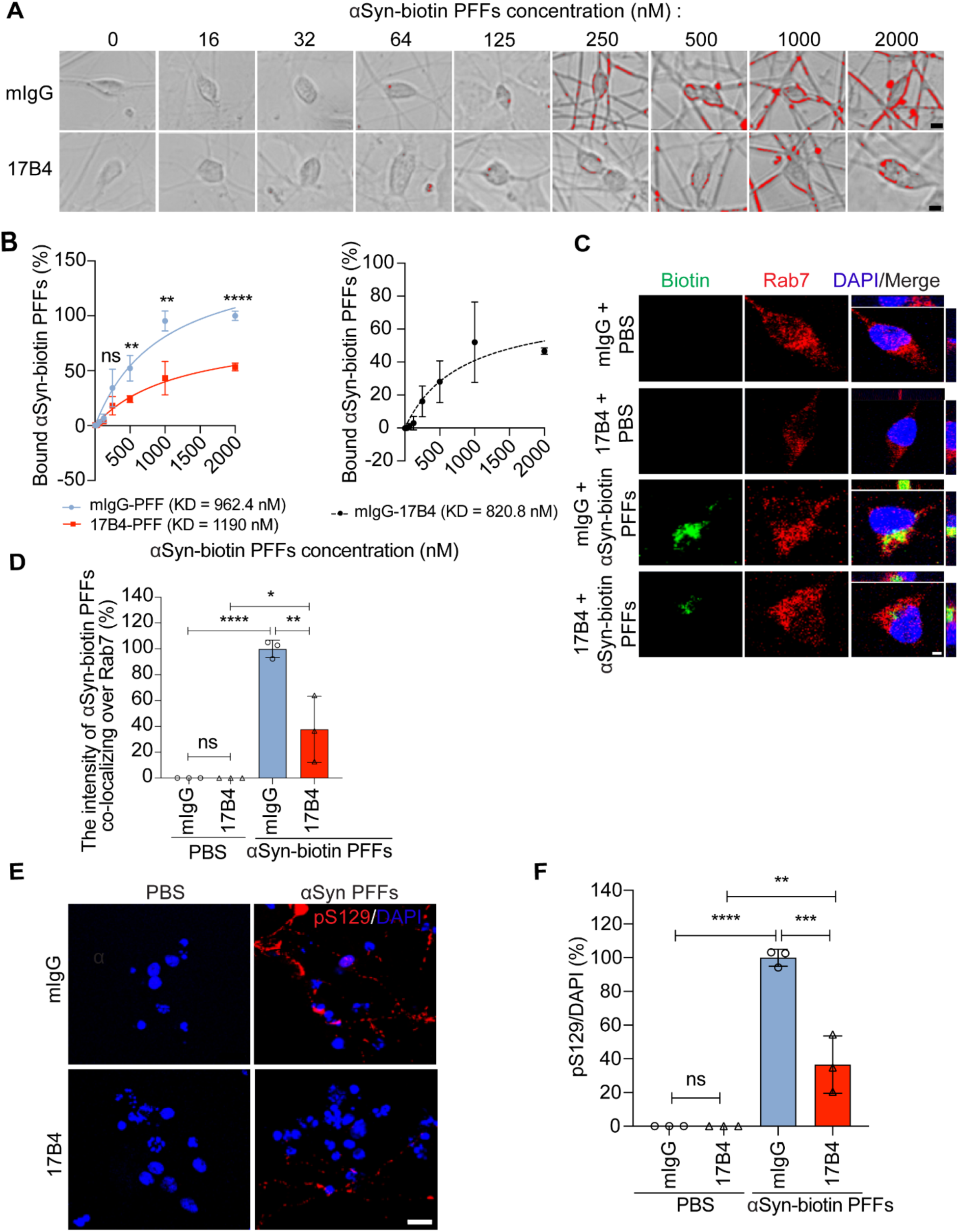
Anti-human LAG3 (17B4) antibody blocks human αSyn PFFs binding, endocytosis, and pathology in induced pluripotent stem cell (iPSC)-derived DA neurons. **(A)** αSyn-biotin PFFs binding to iPSC-derived DA neurons with mouse immunoglobulin G (mIgG) or 17B4 antibody treatment. Binding signal (red) quantified for individual cells. *n* = 9 cells for each group. Scale bar, 5 μm. **(B)** Quantification from *n* = 3 independent experiments. Statistical significance determined by Student’s *t*-test. **(C)** mIgG and 17B4 antibodies blocking human LAG3 inhibit αSyn-biotin PFFs binding on iPSC-derived DA neurons. Colocalization of internalized αSyn-biotin PFFs with Rab7 assessed by confocal microscopy. Scale bar, 5 μm. **(D)** Quantification of colocalization from *n* = 3 independent experiments. **(E)** pS129 signal reduced by 17B4 antibodies in iPSC-derived DA neurons 2 weeks after αSyn PFFs treatment. Scale bar, 10 μm. **(F)** Quantification of pS129 levels from *n* = 3 independent experiments. Statistical significance determined by ANOVA with Tukey’s post-hoc correction. Data presented as means ± SEM. \*\**p* < 0.01, \*\*\**p* < 0.001, \*\*\*\**p* < 0.0001, ns, not significant.

### Anti-Human LAG3 Antibody Inhibits Neuronal Uptake of αSyn PFFs and Suppresses αSyn PFFs-Induced Pathology Propagation in Human DA Neurons

To investigate whether LAG3 plays a role in the endocytosis of αSyn PFFs, we used Rab7, a well-established marker of endocytosis^10, 12, 22^, to assess co-localization with αSyn-biotin PFFs. Human iPSC-derived DA neuron cultures were pretreated with anti-human LAG3 antibody (17B4) or control mouse IgG (mIgG) for two hours, followed by the administration of αSyn-biotin PFFs. In mIgG-treated human DA neurons, a substantial amount of αSyn-biotin PFFs co-localized with Rab7, compared to PBS-treated negative controls (Fig. 4C, D, Fig. S4C). Treatment with the anti-human LAG3 antibody 17B4 significantly reduced the co-localization of αSyn-biotin PFFs with Rab7 compared to mIgG-treated neurons, indicating reduced neuronal uptake by 17B4 (Fig. 4C, D, Fig. S4C).

To further examine the role of human neuronal LAG3 in mediating αSyn pathology propagation, we pre-treated human DA neurons with 17B4 or mIgG two hours before administering αSyn PFFs. Two weeks post-treatment, we observed robust anti-pS129 immunoreactivity in αSyn PFFs-treated neurons compared to PBS-treated controls (Fig. 4E, F). Importantly, pre-treatment with 17B4 significantly suppressed αSyn PFFs-induced pS129 immunoreactivity in human DA neurons compared to the mIgG-αSyn PFFs group (Fig. 4E, F).

### Discussion and Conclusion

In this study, we investigated the neuronal role of LAG3 in mediating αSyn PFFs binding, uptake, and propagation *in vitro* and *in vivo*, as well as its impact on neuronal function ex vivo, and neurodegeneration and behavioral deficits *in vivo*. Our findings demonstrate that conditional knockout of neuronal LAG3 significantly reduces αSyn pathology transmission, alleviates motor dysfunction, and inhibits neurodegeneration induced by αSyn PFFs *in vivo*. Genetic deletion of LAG3 in neurons effectively mitigates the functional deficits induced by αSyn PFFs. Furthermore, treatment with an anti-human LAG3 antibody inhibits the binding and uptake of αSyn PFFs by human DA neurons and suppresses αSyn PFFs-induced pathology propagation in these neurons.

A key question is whether LAG3 expression is in neurons. One prior publication^14^ claimed that LAG3 is not expressed in neurons, but this conclusion was based on the use of an inappropriate positive control—T cells, which express very high LAG3 levels— rather than using LAG3 deficient neurons as the negative control. In contrast, our recent work has confirmed LAG3 expression in neurons using several experimental approaches^12^. Immunostaining in various mouse brain regions, including the cortex, midbrain, thalamus, and hippocampus, demonstrated co-localization of LAG3 with NeuN, a neuronal marker, with approximately 50% of neurons identified as LAG3- positive^12^. RNAscope in situ hybridization^12^ revealed detectable LAG3 mRNA in neurons, particularly in the ventral midbrain and cortex, with no signal observed in LAG3^-/-^ mice^12^. Data from public databases, such as the Allen Brain Atlas and EMBL-EBI Expression Atlas also revealed detectable LAG3 mRNA in neurons^15–21^. In LAG3^L/L-YFP^ reporter mice, YFP immunoreactivity was observed in NeuN-positive and MAP2- positive neurons in the cortex, further supporting the presence of LAG3 in neurons^12^.

Additionally, primary cortical neuron cultures from WT mice showed LAG3 protein expression, while this was absent in LAG3^-/-^ neurons, validating the specificity of the findings^12^.

In this work, we utilized neuronal LAG3 conditional knockout mice and human iPSC-derived DA neurons to confirm the role of neuronal LAG3 in facilitating pathogenic αSyn propagation. The facilitating role of LAG3 in αSyn propagation has also been independently validated in studies of microvascular endothelial cells^23^. Beyond the αSyn PFFs model, we also used transgenic human A53T mutant αSyn mice (hA53T, G2-3 line) to study the role of LAG3^9^. Depletion of LAG3 in these mice significantly prolonged their survival, alleviated behavioral deficits, and inhibited pS129-positive αSyn pathology *in vivo*^9^. These results collectively confirm that LAG3 is expressed in neurons and underscore its critical role in facilitating the progression of α-synucleinopathies.

Another important question is whether LAG3 specifically binds to α-syn fibrils, but not to other prion-like fibrils. We have confirmed that LAG3 does not bind to heparin-induced tau fibrils, as demonstrated in previous studies^10, 11, 22^. However, LAG3 binds strongly to tau fibrils generated from postmortem brain extracts of AD patients^22^. Additionally, we have shown that neuronal LAG3 facilitates pathogenic tau propagation using neuronal conditioned knockout mice.

PD and other prion-like neurodegenerative disorders are driven in part by the spread of pathogenic αSyn and tau, which may be major contributors to disease pathogenesis.

Additional research is required to fully understand how αSyn and tau are transmitted from cell to cell, using appropriate negative controls, disease models, and sufficient group sizes. Further investigation into the interplay between LAG3 and prion-like seeds in aging-related neurodegenerative disorders holds particular promise for uncovering the underlying mechanisms of spread of pathologic proteins.

## Materials and Methods

### αSyn Purification and PFFs and PFFs-Biotin Preparation

Recombinant human αSyn proteins were isolated following established protocols^27^. Human αSyn PFFs were generated by stirring αSyn in a transparent glass vial with a magnetic stirrer at 350 rpm and 37°C. After a 7-day incubation period, the resulting αSyn aggregates were sonicated for 30 seconds at 10% amplitude using a Branson Digital Sonifier (Danbury, CT, USA). The αSyn monomer and PFFs were then separated through Fast Protein Liquid Chromatography (FPLC) on a Superose 6 10/300GL column (GE Healthcare, Life Sciences, Wauwatosa, WI, USA), and the fractions containing human αSyn monomer and PFFs were stored at -80°C. To analyze αSyn PFFs mediators, recombinant αSyn monomer was purified and conjugated with sulfo-NHS-LC-Biotin (Thermo Scientific, Grand Island, NY, USA; EZ-link Sulfo-NHS-LC-Biotin, catalog number 21435) at a biotin-to-αSyn molar ratio of 2 to 3. Purified αSyn underwent endotoxin removal using High-Capacity Endotoxin Removal Spin Columns (Pierce, Rockford, IL, USA) and Ni Sepharose 6 Fast Flow resin (GE Healthcare). The effectiveness of endotoxin removal was then verified with the LAL Chromogenic Endotoxin Quantitation Kit (Pierce).

### Transmission Electron Microscopy (TEM) Measurements

Protein samples (approximately 100 ng/μL) were adsorbed onto 400-mesh carbon-coated copper grids for 5 minutes. The grids were rinsed three times with 50 mM Tris-HCl buffer (pH 7.4) to remove unbound material. Next, the grids were floated on two successive drops of 2% uranyl formate for staining. Excess stain was blotted away with filter paper, and the grids were air-dried before imaging with a Phillips CM 120 TEM.

### Mouse Strain

The LAG3^L/L-YFP^ was obtained from Dr. Dario Vignali at the University of Pittsburgh. Nestin^Cre^ mice were obtained from the Jackson Laboratory (strain #: 003771). LAG3 neuronal conditional knockout (LAG3^L/L-N-/-)^ mice were generated by breeding LAG3^L/L-YFP^ mice with Nestin^Cre^ mice. These mice exhibit no evidence of autoimmune or inflammatory phenotypes. C57BL/6WT (Strain #:000664, RRID: IMSR JAX:000664) were obtained from the Jackson Laboratories (Bar Harbor, ME). LAG3^-/-^ mice^38^ were obtained from Dr. Charles G. Drake when he was at Johns Hopkins University. Animals were housed in ventilated cages with unlimited access to food and water. All the procedures were approved by the Johns Hopkins Animal Care and Use Committee and carried out following the guidelines from NIH for Care and Use of Experimental Animals.

### Stereotaxic Injection of αSyn PFFs and Sample Preparation

On the day of intrastriatal injections, αSyn PFFs was diluted in sterile PBS and briefly sonicated. Three-month-old mice were anesthetized with a mixture of ketamine (100 mg/kg) and xylazine (10 mg/kg), followed by intrastriatal injection of αSyn PFFs (5 µg/2 µL at 0.4 µL/min). The injection targeted the dorsal striatum using the following coordinates from bregma: anteroposterior (AP) = +0.2 mm, mediolateral (ML) = +2.0 mm, dorsoventral (DV) = +2.8 mm with a 2 µL syringe (Hamilton, USA). The needle was kept in place for an additional 5 minutes to ensure complete diffusion before slow withdrawal. Post-surgery, animals were monitored, and all received appropriate care. Behavioral tests were conducted 6 months post-αSyn PFFs injection. Mice were euthanized at two or three months after surgery for neurochemical, biochemical, and histological analyses. Tissues for biochemical experiments were immediately removed, snap-frozen, and stored at −80°C. For histological study, mice were perfused transcardially with PBS, followed by fixation with 4% paraformaldehyde (PFA). The brains were then extracted, post-fixed overnight in 4% PFA, and subsequently transferred to 30% sucrose in PBS for cryoprotection.

### Immunofluorescence and Mapping of pS129-Positive αSyn Pathology *In Vivo*

Coronal brain sections (30 µm thick) were incubated with primary antibodies against phospho-serine 129 αSyn (ab51253, Abcam) for mapping in various brain regions. Additional antibodies used included NeuN (ab134014, Abcam) and LAG3 (LS-B15026, LSBio) on tissue sections, followed by matched fluorescence-conjugated secondary antibodies (Invitrogen). Nuclei were counterstained with DAPI. Images were captured using confocal scanning microscopy (LSM 980, Zeiss, Dublin, CA, USA) or a fluorescent microscope (BZ-X710, Keyence, Osaka, Japan).

### Immunohistochemistry and Stereotaxic Counting of Tyrosine Hydroxylase and Nissl Positive Cells

Immunohistochemistry was conducted on 30 µm thick serial brain sections. Free-floating sections were blocked with 4% goat serum (Sigma-Aldrich) in PBS containing 0.3% Triton X-100, followed by incubation with primary antibodies against tyrosine hydroxylase (TH). Secondary antibody incubation was performed using biotin-conjugated anti-rabbit antibodies. Antibody-antigen complexes were visualized using ABC reagents (Vector Laboratories) and Sigma Fast DAB Peroxidase Substrate (Sigma-Aldrich). TH-stained sections were counterstained with Nissl (0.09% thionin) as previously described. Sections were then dehydrated in 100% ethanol, cleared in xylene (Fisher Scientific), and mounted with DPX (Sigma-Aldrich) before imaging under a microscope. Cell counting was conducted using an unbiased stereological method with the optical fractionator approach. Counts of TH-positive or Nissl-positive neurons were performed in the substantia nigra of the right hemisphere of mouse brains. The assessments were conducted by experimenters blinded to both genotype and treatment groups. Stereological analysis utilized a computer-assisted image analysis system comprising an Axiophot photomicroscope (Carl Zeiss Vision), a motorized stage (Ludl Electronics), a Hitachi HV C20 video camera, and Stereo Investigator software (MicroBrightField). Total counts of TH- and Nissl-positive neurons were calculated as previously described^39, 40^.

### Tissue Lysate Preparation and Immunoblot Analysis

Tissue lysates were prepared for immunoblot analysis. Dissected brain regions were homogenized in RIPA buffer supplemented with Protease/Phosphatase Inhibitor Cocktail (5872S, Cell Signaling Technology, Danvers, MA). Following centrifugation at 10,000 rpm for 20 minutes, the supernatants were collected, and protein concentrations were quantified using the BCA assay (Pierce, USA). Samples containing 20-30 µg of total protein were separated on 12.5–13.5% SDS-polyacrylamide gels, then transferred onto PVDF membranes and blocked with 5% nonfat milk or 5% BSA in TBS-T (Tris-buffered saline, 0.1% Tween 20). Membranes were incubated with primary and secondary antibodies, signals were detected using ECL or SuperSignal Femto substrate (Thermo Fisher, USA), and images were captured with an ImageQuant LAS 4000 mini scanner (GE Healthcare Life Sciences, USA).

### Behavioral analysis

Behavioral analyses were performed 10 days prior to the euthanasia of mice to assess behavioral deficits induced by PFFs as well as the negative control PBS. Tests conducted included the pole test, grip strength, rotarod test, and nest building, all of which were administered by experimenters who were blinded to the treatments and randomly assigned to groups.

### Pole Test

Mice were acclimated to the behavioral procedure room for 30 minutes before testing. The apparatus was a 75-cm long metal rod with a 9 mm diameter, wrapped in bandage gauze for grip. For the test, mice were placed near the top of the pole (7.5 cm from the top) in an upward-facing orientation. The total time for each mouse to descend to the base of the pole was recorded. To familiarize the mice with the task, they underwent training for two consecutive days, with each session consisting of three test trials. On the test day, performance was evaluated over three sessions, recording the total time. A maximum cutoff time of 60 seconds was established for each trial, after which the test was terminated, and the time was recorded. Outcomes measured included turn-down time, climb-down time, and total time (in seconds).

### Grip Strength

Neuromuscular strength was assessed by measuring the maximum holding force of mice with a force transducer (Bioseb, USA). Mice were allowed to grip a metal grid using their forelimbs, hind limbs, or both, while their tails were gently pulled. The peak force at the moment the grip was lost was recorded by the transducer and quantified in grams.

### Rotarod Test

Mice were conditioned for three consecutive days before testing to familiarize them with the task. On the test day, they were placed on an accelerating rotarod where the latency to fall was recorded. The rotarod’s speed increased from 4 to 40 rpm over 5 minutes. A trial ended if a mouse fell or if it gripped the rod and spun for two consecutive cycles without attempting to walk. Motor performance was reported as the percentage of mean latency to fall (averaged from three trials) relative to the control group.

### Nest Building

The nest-building test was used to assess nigrostriatal sensorimotor and hippocampal cognitive function. This test involves orofacial and forelimb movements where mice manipulate nesting material with their forelimbs and teeth, breaking it down and integrating it into their bedding. For this study, a 2.5 g nestlet (Johns Hopkins Medicine Research Animal Resources, Baltimore, MD, USA) was placed in each cage’s feeder, requiring the mice to perform complex fine motor tasks to retrieve the material. After 16 hours, nest-building was scored, and the amount of unused nesting material was measured as an indicator of sensorimotor function.

### Electrophysiology methods

#### Slice preparation

Mice were anesthetized with Euthasol and decapitated, brains were quickly removed and placed in ice-cold low-sodium ACSF. Horizontal midbrain slices (200 – 250 μm) containing the SNc were prepared in ice-cold ACSF using a vibrating blade microtome (Leica VT1200). Right after cutting, slices were recovered for 10 minutes at 32°C degrees and then transferred to holding ACSF at room temperature. Cutting and recovery were performed with ACSF containing the sodium substitute NMDG^41^ (in mM): 92 NMDG, 20 HEPES (pH 7.35), 25 glucose, 30 sodium bicarbonate, 1.2 sodium phosphate, 2.5 potassium chloride, 5 sodium ascorbate, 3 sodium pyruvate, 2 thiourea, 10 magnesium sulfate, 0.5 calcium chloride. ACSF used for holding slices prior to recording was identical, but contained 92 mM sodium chloride, instead of NMDG, 1 mM magnesium chloride and 2 mM calcium chloride. ACSF used to perfuse slices during recording contained (in mM): 125 sodium chloride, 2.5 potassium chloride, 1.25 sodium phosphate, 1 magnesium chloride, 2.4 calcium chloride, 26 sodium bicarbonate, and 11 glucose. All ACSF solutions were saturated with 95% O_2_ and 5% CO_2_. After recovering slices were incubated with αSyn-PFFs, and PBS (used as control condition) for 90 min at room temperature in ACSF. The length of the incubation protocol is in accordance with previous studies^31–33^ and relates to the main purpose of the present study, which was to evaluate the impact of an acute exposure of midbrain DA neurons to extracellular form of αSyn on the intrinsic excitability properties and the synaptic transmission. The range of concentration and time-course chosen for αSyn PFFs incubation were tested and optimized for obtaining the max effect without compromising the slice viability crucial for the electrophysiology recordings. All the experiments were carried out at the 10 μg/ml (700 nM) of αSyn PFFs. Following this incubation period, slices were then washed, placed in the recording chamber and perfused with ACSF at 32°C. The experimental procedure is summarized in Figure 3B. The control condition consisted of PBS incubation at same volume used for αSyn PFFs.

#### Whole-cell and cell-attached patch-clamp electrophysiology

Horizontal midbrain slices containing were transferred to a small volume (0.5 ml) recording chamber, mounted on a fixed stage upright microscope equipped with IR-DIC (BX51, Olympus America). All the experiments were performed at a temperature ranging from 31 to 33°C, unless specified otherwise. The recording chamber was continuously perfused with carbogen-saturated ACSF at a flow rate of 2–2.5 ml/min using a peristaltic pump (WPI) running through an in-line heater (SH-27B with TC-324B controller; Warner Instruments).

Following previous studies, the medial terminal (mt) nucleus of the accessory optic tract serves as useful landmark to localize SNc, respectively. The SNc DA neurons were defined in the region laterally to the *mt* nucleus, between the medial lemniscus dorsally and the substantia nigra pars reticulata ventrally^42, 43^. This region contains a dense band of neurons with large cell bodies and with dendritic processes running in a mediolateral manner when visualized under IR-DIC microscopy. Conventional tight-seal whole-cell patch- clamp and cell-attached recordings were made on visually identified, DA neurons, based on size and morphology. DA neurons were also identified by their physiological features, including resting level of discharge, presence of a prominent voltage sag during hyperpolarizing current injection and high spike threshold (typically above 240 mV), which are considered reasonable predictors of dopaminergic identity in mice^44, 45^. Neurobiotin (0.05%; Vector Labs, Burlingame, CA) was included in the intracellular solutions to allow identification of dopaminergic neurons using post-hoc tyrosine-hydroxylase immunolabeling^46^. Recordings were made using a MultiClamp 700B amplifier (1 kHz low-pass Bessel filter and 10 kHz digitization) with pClamp 10.3 software (Molecular Devices). Patch electrodes (1.5 mm outer diameter) were fabricated from filamented, thick-wall borosilicate glass pulled on a Flaming-Brown puller (P-85, Sutter Instruments) and fire-polished immediately before use. Glass pipets typical resistance was 1.8-3 MΩ, when filled with internal solutions. For *spontaneous and evoked firing,* the recording internal solution consisted of (mM): 135 K-gluconate, 4 KCl, 10 HEPES, 4 Mg-ATP, and 0.3 Na-GTP, adjusting with KOH for a final pH 7.2–7.3 and 280–285 mOsm. To avoid contamination from fast excitatory and inhibitory synaptic transmission DNQX (10μM; Tocris), D-APV (50μM; Tocris) and picrotoxin (100μM; ABCAM) were added to the extracellular ACSF solution. To maximize the number of recordings displaying healthy firing activity, spontaneous firing was first monitored for a period in a cell-attached configuration. The membrane was then ruptured while recording in current-clamp mode, which allowed for easy identification of firing rate changes due to damage during breakthrough. Initial spontaneous firing activity was obtained for at least 5 min, during which the health of the neuron was further evaluated. Only cells that were spontaneously active after breaking through the whole-cell configuration were included in the analysis. Frequency-intensity curves were obtained for each neuron by providing a series of depolarizing current injections (50 pA steps, 2 s duration) during their spontaneous activity. Input and series resistance were continuously monitored on-line; if either parameter changed by more than 20% data were not included in the analysis. Membrane potentials were not corrected for junction potentials (estimated to be 10 mV). A bipolar stimulating electrode was placed rostrally at a distance of 100–300 μm from the recording electrode with a mediolateral position optimal for evoking synaptic activity. The internal recording solution was composed of (in mM): 117 cesium methanesulfonate, 20 HEPES, 0.4 EGTA, 2.8 NaCl, 5 TEA-Cl, 2.5 Mg-ATP, and 0.25 Na-GTP, pH 7.2–7.3 and 280–285 mOsm. Picrotoxin (100 μM) was added to block GABAA receptor-mediated inhibitory postsynaptic currents. mEPSCs and mIPSCs were pharmacologically isolated by having picrotoxin (100 μM) plus tetrodotoxin (1 μM), or DNQX (10 μM) plus D-APV (50 μM) and tetrodotoxin (1 μM), present throughout the experiment, and sampled at 1 KHz while clamping the cells at −70 mV and 0 mV, respectively. 150 events per cell were acquired and detected using a threshold of 8 pA. Analysis of mEPSCs and mIPSCs were performed off-line and verified by eye using the MiniAnalysis program (v 6.0, Synaptosoft).

#### Immunohistochemistry

Acute slices containing Neurobiotin tracer-labeled cells were fixed for 60 min in 4% paraformaldehyde at 4°C and immunolabelled with anti-tyrosine hydroxylase (Sigma Aldrich, 1:500) and Streptavidin Alexa Fluor 594 (Invitrogen; 1:12,000) and donkey anti-sheep Alexa Fluor 488 (Invitrogen; 1:1000 2 μg/ml) as previously described. All immunolabelling was viewed on an Olympus confocal microscope (Leica Microsystems, Wetzlar, Germany), and images were captured using Olympus software. Figures were prepared using Corel Draw and Adobe Illustrator (CS4, Adobe Systems Inc., San Jose, CA, U.S.A.) and ImageJ (MacBiophotonics, McMaster University, ON, Canada), with brightness and contrast adjustments performed consistently across the image to enhance clarity. *<u>Drugs</u>*. picrotoxin, D-APV, ifenprodil were from ABCAM; all the others from Tocris. *<u>Statistical analysis.</u>* Electrophysiological data collected were analyzed using ClampFit 10.2 (Molecular Devices), MiniAnalysis program (v 6.0, Synaptosoft). Statistical analysis was done with Graphpad Prism 7.03. Parametric and non-parametric testing were appropriately chosen after performing normality test. The values are presented as mean ± SEM. A probability value of *p* < 0.05 was considered statistically significant (\**p* < 0.05; \*\**p* < 0.01; \*\*\**p* < 0.001; \*\*\*\**p* < 0.0001). In statistical tests between 3 or more conditions, a one-way ANOVA followed by a Bonferroni’s multiple comparison post *hoc* test was used.

### Generation and Differentiation of DA-Neuronal Cells from Atoh1-Transduced iPSCs for αSyn PFFs Studies

Atoh1-transduced iPSCs were utilized to generate DA-neuronal cells^35^. iPSCs were cultured under feeder-free conditions in mTeSR™ Plus medium (StemCell Technologies; 100-0276) within a humidified incubator at 37°C with 5% CO_2_/95% air. For differentiation, cells were seeded at a density of 8 × 10⁴ cells/cm² on Matrigel-coated plates in mTeSR™ Plus medium containing the ROCK inhibitor Y-27632 (10 μM; Stemgent). Doxycycline (0.5 μg/mL) was added to the culture medium from day 1 to day 5 to induce Atoh1 expression. The medium was replaced daily from day 1 to day 3, transitioning gradually from mTeSR™ Plus to N2 medium (DMEM/F-12 with N2 supplement; Life Technologies). Cells were maintained in N2 medium until day 5, then dissociated with Accutase (Sigma-Aldrich) and replated at a density of 3 × 10⁵ cells/cm² on poly-D-lysine (1 μg/mL) and laminin (1 μg/mL)-coated plates in neuron culture medium. The neuron medium consisted of Neurobasal medium supplemented with B27, BDNF (10 ng/mL), GDNF (10 ng/mL), TGFβ-3 (1 ng/mL), cAMP (0.1 mM), ascorbic acid (0.2 mM), and DAPT (10 μM). Medium replenishment occurred every 3–4 days. By day 7 of differentiation, Atoh1-induced DA neuron precursors were collected using Accutase for further applications. For all experiments, DA-neuronal cells were pretreated with either mIgG or 17B4 (mouse anti-human LAG3, LifeSpan Biosciences, LS-C344746) for 2 hours. Subsequently, human αSyn-biotin PFFs were applied for 1.5 hours. In pathology-focused studies, human αSyn PFFs exposure was extended to a total duration of 2 weeks.

### Immunofluorescence and Mapping of pS129-Positive αSyn Pathology *In Vitro*

For pathology studies in iPSC-derived DA-neuronal cells, cells were pretreated with either mIgG or 17B4 (5 μg/mL) for 2 hours. Following pretreatment, PBS or human αSyn PFFs (5 μg/mL) were applied. Medium was replenished every 4 days by replacing half with fresh neuron culture medium. After 2 weeks, cells were fixed with 4% paraformaldehyde/1% sucrose in PBS. Blocking was performed using 10% horse serum and 0.1% Triton X-100, followed by incubation with the pS129 antibody (Abcam, ab51253; 1:1000). Immunoreactivity was visualized using Alexa Fluor 568-conjugated secondary antibodies (Invitrogen, Carlsbad, CA, USA). Nuclei were counterstained with DAPI alongside the secondary antibodies. Images were acquired using a confocal scanning microscope (LSM 880, Zeiss, Dublin, CA, USA) or a fluorescence microscope (BZ-X710, Keyence, Osaka, Japan).

### Cell Surface Binding Assays for αSyn PFFs

iPSC-derived DA-neuronal cells were treated with mIgG or 17B4 (5 μg/mL) antibody for 2 hours at room temperature (RT). Following this pretreatment, the cells were incubated with varying concentrations of human αSyn-biotin PFFs, up to 2000 nM, in neuron culture medium for 1.5 hours at RT^10–12, 22^. Unbound αSyn-biotin PFFs were removed through five thorough washes (20 minutes each) with neuron culture medium on a horizontal shaker. Cells were subsequently fixed with 4% paraformaldehyde/1% sucrose in PBS and rinsed three times with PBS. Blocking was performed for 30 minutes using PBS containing 10% horse serum and 0.1% Triton X-100. After blocking, cells were incubated for 16 hours with alkaline phosphatase-conjugated streptavidin (1:2000 dilution) in PBS containing 5% horse serum and 0.05% Triton X-100. Alkaline phosphatase activity was visualized histochemically using a BCIP/NBT reaction. Microscopic images were captured using a Zeiss Axiovert 200M microscope. Quantification of αSyn-biotin PFFs binding intensity to iPSC-derived DA-neuronal cells was performed using ImageJ software. Thresholds were adjusted under the Image/Adjust menu to exclude background signals, and identical parameters were applied across all images within each experiment. GraphPad Prism software was used to calculate binding curves and dissociation constant (KD) values.

### ImageJ Analysis

The threshold was adjusted in Image/Adjust to define an optimal intensity range for each experiment. This threshold was applied consistently across all images within the same experiment to ensure uniform analysis. The threshold setting also facilitated the exclusion of background noise. After background removal, regions within the defined signal intensity range were quantified using the measurement tool in the Analyze/Measure menu. The measured area values corresponding to varying concentrations of αSyn-biotin (monomer or PFFs) were subsequently analyzed using Prism software to calculate the dissociation constant (KD).

### Colocalization of Rab7 and αSyn-Biotin PFFs

iPSC-derived DA-neuronal cells were pretreated with mIgG or 17B4 (5 μg/mL) for 2 hours. Following pretreatment, the cells were incubated with PBS or human αSyn-biotin PFFs (5 μg/mL) for 1.5 hours. After three PBS washes, the cells were fixed with 4% paraformaldehyde/1% sucrose in PBS. Post-fixation, cells were washed three additional times with PBS and blocked for 1 hour in PBS containing 10% horse serum and 0.1% Triton X-100. After blocking, cells were stained with Rab7 (1:1000) and streptavidin-conjugated Alexa Fluor 647 (1:1000). Images were captured under identical exposure settings and processed uniformly for analysis. The colocalization of αSyn-biotin PFFs with Rab7 was evaluated and quantified using Zeiss Zen software. Regions of colocalization were outlined, and the intensity and area of the colocalized signals were measured. Total signal values were calculated by multiplying the measured intensity by the area to determine the overall colocalization value.

## Acknowledgements

The authors acknowledge the joint participation by the Adrienne Helis Malvin Medical Research Foundation through its direct engagement in the continuous active conduct of medical research in conjunction with The Johns Hopkins Hospital and the Johns Hopkins University School of Medicine and the Foundation’s Parkinson’s Disease Program M- 2023. TMD is the Leonard and Madlyn Abramson Professor in Neurodegenerative Diseases. The Multiphoton Imaging Core of Johns Hopkins University was used (NS050274) in some of the imaging studies.

## Author contributions

XM designed and led the project and contributed to all aspects of the study; XLY, DJ, GM contributed to all the experiments, data analysis, and interpretation; RK, NW, FA contributed to biochemical, cellular, mouse experiments; CJW, DAAV provided key tools, reagents or critical information or dataset; MYY helped designed research and data interpretation; MYY, VLD, TMD, XM wrote the paper. All authors reviewed, edited, and approved the paper.

## Data and materials availability

Further information and requests for resources and reagents should be directed to and will be fulfilled by the Lead Contact, Xiaobo Mao. There are no restrictions on any data or materials presented in this paper. All data are available in the main text or the Extended Data.

---

**Fig. S1.**
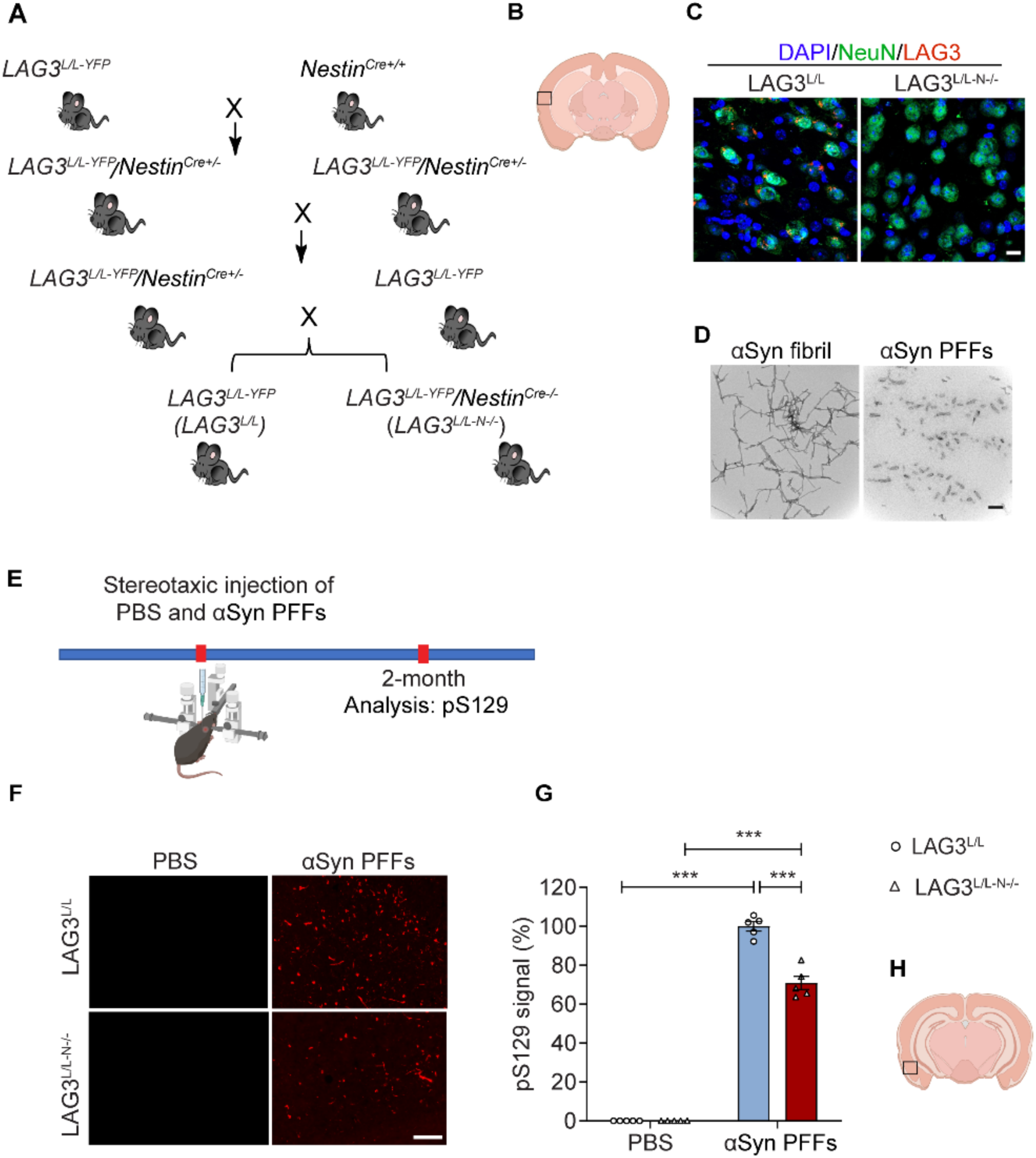
Attenuation of αSyn PFFs-induced pS129-positive pathology in neuronal LAG3 conditional knockout (LAG3^L/L-N-/-^) mice. **(A)** Breeding strategy for generating LAG3^L/L-N-/-^ mice. **(B, C)** Confirmation of LAG3 deletion from neurons in LAG3^L/L-N-/-^ mice. **(D)** Transmission electron microscopy (TEM) images of αSyn PFFs. Scale bar, 100 nm. **(E—G)** Immunohistochemical staining of pS129 in the entorhinal cortex at 3 months post- αSyn PFFs injection. Quantification from *n* = 5 biologically independent experiments. Statistical significance determined by ANOVA with Tukey’s post-hoc correction. Data presented as means ± SEM. \*\*\**p* < 0.001.

**Fig. S2.**
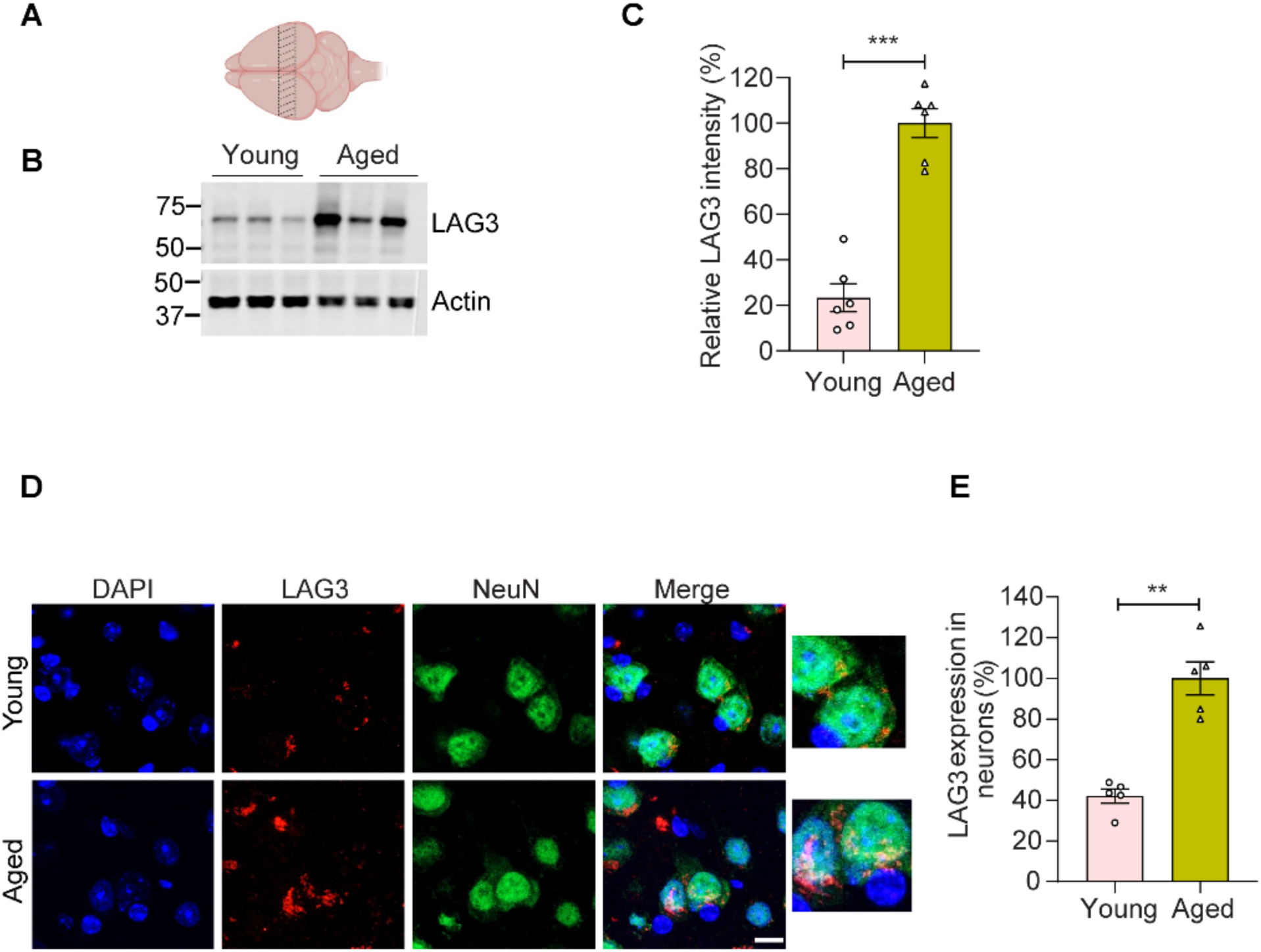
Decreased expression of LAG3 in young mice compared to aged mice. **(A)** Representative microphotographs showed midbrain tissue for protein lysis. **(B)** Representative immunoblots of LAG3 and β-actin in the ventral midbrain of young and aged mice. **(C)** Quantification of LAG3 levels. Data are mean ± SEM; *n* = 6 biologically independent animals. *\*\*\*p* < 0.001. **(D, E)** Representative images, and percentage of Lag3 positive in the neuron of young and aged mice. Scale bar, 10 μm. Data are mean ± SEM, statistical significance was calculated with unpaired two tailed Student’s *t*-tests. *n* = 5 biologically independent animals. *\*\*p* < 0.01

**Fig. S3.**
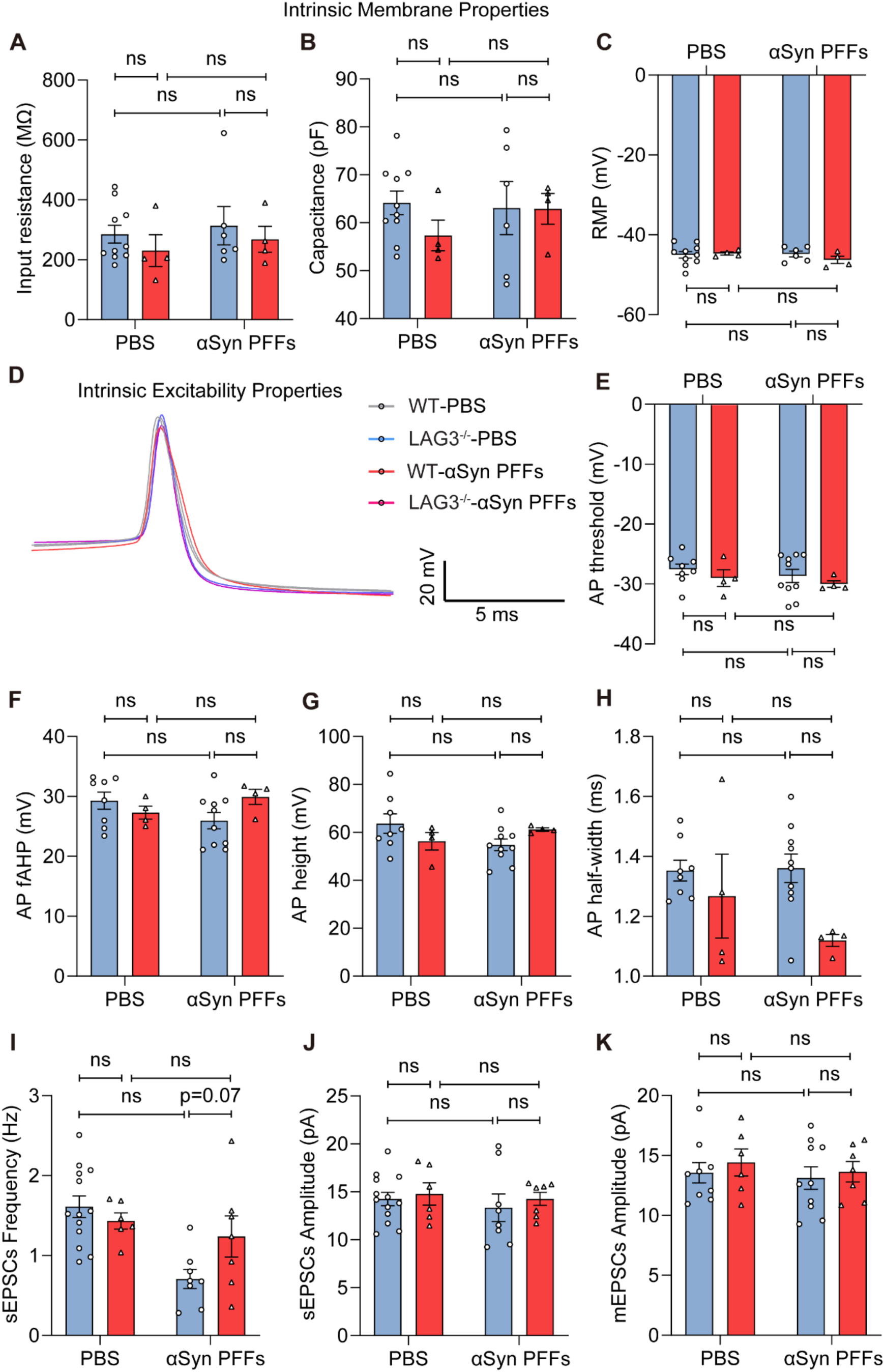
No changes in specific intrinsic properties of SNc DA neurons following *in vitro* incubation with αSyn PFFs in WT and LAG3^-/-^ mice. (A—C) Intrinsic membrane properties of SNc DA neurons: **(A)** Input resistance, **(B)** Capacitance, **(C)** Resting membrane potential (RMP) in αSyn PFFs and PBS conditions for WT and LAG3^-/-^ mice. SNc DA neurons from WT and LAG3^-/-^ mice show no significant changes in intrinsic membrane properties following 90-minute incubation with αSyn PFFs or PBS. WT mice- PBS, *n* = 10; LAG3^-/-^-PBS mice, *n* = 4; WT mice-αSyn PFFs, *n* = 6; LAG3^-/-^ mice-αSyn PFFs, *n* = 4. **(D)** Normalized average action potential (AP) waveform (grey = WT mice- PBS, blue = LAG3^-/-^ -PBS mice, red = WT mice-αSyn PFFs, purple = LAG3^-/-^ mice-αSyn PFFs). **(E—H)** No significant changes in AP parameters: **(E)** AP threshold, **(F)** Fast afterhyperpolarization (fAHP), **(G)** AP height, **(H)** AP half-width in SNc DA neurons of both WT and LAG3^-/-^ mice after 90-minute αSyn PFFs incubation compared to PBS incubation. WT mice-PBS, *n* = 8; LAG3^-/-^-PBS mice, n = 5; WT mice-αSyn PFFs, *n* = 10; LAG3-/- mice-αSyn PFFs, *n* = 4. **(I)** Spontaneous excitatory postsynaptic currents (EPSC) frequency in SNc DA neurons from PBS and αSyn PFFs-treated WT and LAG3^-/-^ mice using whole-cell recording. **(J)** Spontaneous excitatory postsynaptic current (EPSC) amplitude in SNc DA neurons from PBS and αSyn PFFs-treated slices of WT and LAG3^-/-^ mice using whole-cell recording. WT mice-PBS, *n* = 13; LAG3^-/-^-PBS mice, *n* = 6; WT mice-αSyn PFFs, *n* = 8; LAG3^-/-^ mice-αSyn PFFs, *n* = 7. **(K)** Miniature EPSC amplitude in SNc DA neurons from PBS and αSyn PFFs-treated slices of WT and LAG3^-/-^ mice using whole-cell recording. WT mice-PBS, *n* = 9; LAG3^-/-^-PBS mice, *n* = 6; WT mice-αSyn PFFs, *n* = 10; LAG3^-/-^ mice-αSyn PFFs, *n* = 7. Values expressed as mean ± SEM. ns, not significant. Statistical significance determined by ANOVA.

**Fig. S4.**
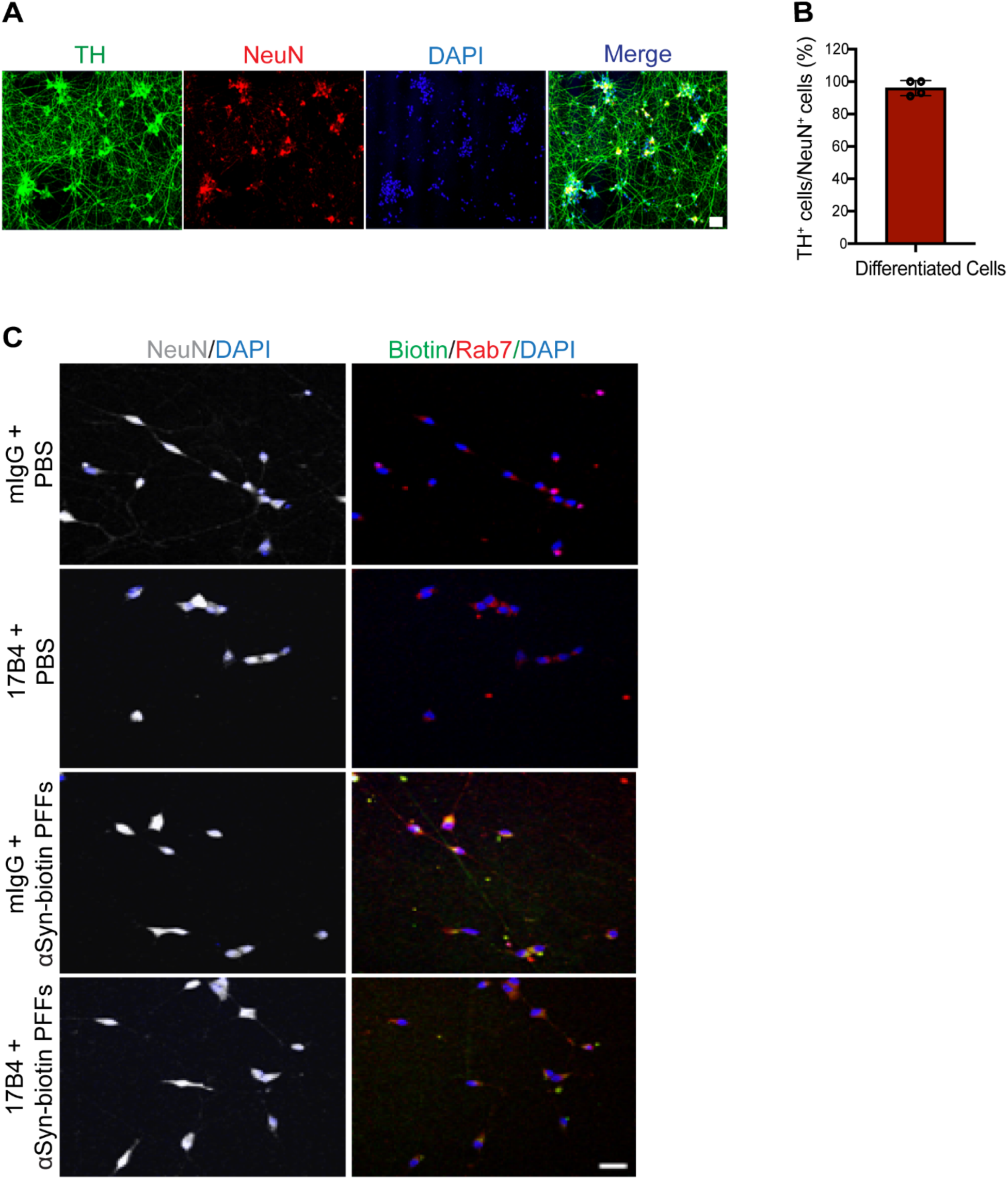
Characterization of human DA neurons derived from induced pluripotent stem cells (iPSCs) and reduction of αSyn PFFs endocytosis by 17B4 antibody. **(A)** Immunofluorescence staining of iPSC-derived DA neurons with tyrosine hydroxylase (TH, green), neuronal nuclei marker (NeuN, red), and nuclear counterstain (DAPI, blue). Scale bar, 50 μm. **(B)** Quantification of TH and NeuN-positive iPSC-derived DA neurons. **(C)** Mouse immunoglobulin G (mIgG) and 17B4 antibodies targeting human LAG3 reduce colocalization of αSyn-biotin PFFs with late endosome marker Rab7. Scale bar, 20 μm.

## References

1. Volpicelli-Daley LA, Luk KC, Patel TP, Tanik SA, Riddle DM, Stieber A, Meaney DF, Trojanowski JQ, Lee VM. Exogenous alpha-synuclein fibrils induce Lewy body pathology leading to synaptic dysfunction and neuron death. Neuron. 2011;72(1):57–71. Epub 2011/10/11. doi: 10.1016/j.neuron.2011.08.033. PubMed PMID: 21982369; PMCID: PMC3204802.

2. Luk KC, Kehm V, Carroll J, Zhang B, O’Brien P, Trojanowski JQ, Lee VM. Pathological alpha-synuclein transmission initiates Parkinson-like neurodegeneration in nontransgenic mice. Science. 2012;338(6109):949-53. doi: 10.1126/science.1227157. PubMed PMID: 23161999; PMCID: 3552321.

3. Holmes BB, DeVos SL, Kfoury N, Li M, Jacks R, Yanamandra K, Ouidja MO, Brodsky FM, Marasa J, Bagchi DP, Kotzbauer PT, Miller TM, Papy-Garcia D, Diamond MI. Heparan sulfate proteoglycans mediate internalization and propagation of specific proteopathic seeds. Proc Natl Acad Sci U S A. 2013;110(33):E3138-47. Epub 20130729. doi: 10.1073/pnas.1301440110. PubMed PMID: 23898162; PMCID: PMC3746848.

4. Kim C, Ho DH, Suk JE, You S, Michael S, Kang J, Joong Lee S, Masliah E, Hwang D, Lee HJ, Lee SJ. Neuron-released oligomeric alpha-synuclein is an endogenous agonist of TLR2 for paracrine activation of microglia. Nat Commun. 2013;4:1562. doi: 10.1038/ncomms2534. PubMed PMID: 23463005; PMCID: PMC4089961.

5. Daniele SG, Beraud D, Davenport C, Cheng K, Yin H, Maguire-Zeiss KA. Activation of MyD88-dependent TLR1/2 signaling by misfolded alpha-synuclein, a protein linked to neurodegenerative disorders. Sci Signal. 2015;8(376):ra45. Epub 20150512. doi: 10.1126/scisignal.2005965. PubMed PMID: 25969543; PMCID: PMC4601639.

6. Shrivastava AN, Redeker V, Fritz N, Pieri L, Almeida LG, Spolidoro M, Liebmann T, Bousset L, Renner M, Lena C, Aperia A, Melki R, Triller A. alpha-synuclein assemblies sequester neuronal alpha3-Na+/K+-ATPase and impair Na+ gradient. EMBO J. 2015;34(19):2408–23. Epub 20150831. doi: 10.15252/embj.201591397. PubMed PMID: 26323479; PMCID: PMC4601662.

7. Choi YR, Cha SH, Kang SJ, Kim JB, Jou I, Park SM. Prion-like Propagation of alpha-Synuclein Is Regulated by the FcgammaRIIB-SHP-1/2 Signaling Pathway in Neurons. Cell Rep. 2018;22(1):136–48. doi: 10.1016/j.celrep.2017.12.009. PubMed PMID: 29298416.

8. Urrea L, Segura-Feliu M, Masuda-Suzukake M, Hervera A, Pedraz L, Garcia Aznar JM, Vila M, Samitier J, Torrents E, Ferrer I, Gavin R, Hagesawa M, Del Rio JA. Involvement of Cellular Prion Protein in alpha-Synuclein Transport in Neurons. Mol Neurobiol. 2018;55(3):1847–60. Epub 20170222. doi: 10.1007/s12035-017-0451-4. PubMed PMID: 28229331; PMCID: PMC5840251.

9. Gu H, Yang X, Mao X, Xu E, Qi C, Wang H, Brahmachari S, York B, Sriparna M, Li A, Chang M, Patel P, Dawson VL, Dawson TM. Lymphocyte Activation Gene 3 (Lag3) Contributes to alpha-Synucleinopathy in alpha-Synuclein Transgenic Mice. Front Cell Neurosci. 2021;15:656426. Epub 20210310. doi: 10.3389/fncel.2021.656426. PubMed PMID: 33776654; PMCID: PMC7987675.

10. Mao X, Ou MT, Karuppagounder SS, Kam TI, Yin X, Xiong Y, Ge P, Umanah GE, Brahmachari S, Shin JH, Kang HC, Zhang J, Xu J, Chen R, Park H, Andrabi SA, Kang SU, Goncalves RA, Liang Y, Zhang S, Qi C, Lam S, Keiler JA, Tyson J, Kim D, Panicker N, Yun SP, Workman CJ, Vignali DA, Dawson VL, Ko HS, Dawson TM. Pathological alpha-synuclein transmission initiated by binding lymphocyte-activation gene 3. Science. 2016;353(6307). doi: 10.1126/science.aah3374. PubMed PMID: 27708076; PMCID: PMC5510615.

11. Zhang S, Liu YQ, Jia C, Lim YJ, Feng G, Xu E, Long H, Kimura Y, Tao Y, Zhao C, Wang C, Liu Z, Hu JJ, Ma MR, Liu Z, Jiang L, Li D, Wang R, Dawson VL, Dawson TM, Li YM, Mao X, Liu C. Mechanistic basis for receptor-mediated pathological alpha-synuclein fibril cell-to-cell transmission in Parkinson’s disease. Proc Natl Acad Sci U S A. 2021;118(26). doi: 10.1073/pnas.2011196118. PubMed PMID: 34172566; PMCID: PMC8256039.

12. Mao X, Gu H, Kim D, Kimura Y, Wang N, Xu E, Kumbhar R, Ming X, Wang H, Chen C, Zhang S, Jia C, Liu Y, Bian H, Karuppagounder SS, Akkentli F, Chen Q, Jia L, Hwang H, Lee SH, Ke X, Chang M, Li A, Yang J, Rastegar C, Sriparna M, Ge P, Brahmachari S, Kim S, Zhang S, Shimoda Y, Saar M, Liu H, Kweon SH, Ying M, Workman CJ, Vignali DAA, Muller UC, Liu C, Ko HS, Dawson VL, Dawson TM. Aplp1 interacts with Lag3 to facilitate transmission of pathologic alpha-synuclein. Nat Commun. 2024;15(1):4663. Epub 20240531. doi: 10.1038/s41467-024-49016-3. PubMed PMID: 38821932; PMCID: PMC11143359.

13. Kam TI, Mao X, Park H, Chou SC, Karuppagounder SS, Umanah GE, Yun SP, Brahmachari S, Panicker N, Chen R, Andrabi SA, Qi C, Poirier GG, Pletnikova O, Troncoso JC, Bekris LM, Leverenz JB, Pantelyat A, Ko HS, Rosenthal LS, Dawson TM, Dawson VL. Poly(ADP-ribose) drives pathologic alpha-synuclein neurodegeneration in Parkinson’s disease. Science. 2018;362(6414). doi: 10.1126/science.aat8407. PubMed PMID: 30385548; PMCID: PMC6431793.

14. Emmenegger M, De Cecco E, Hruska-Plochan M, Eninger T, Schneider MM, Barth M, Tantardini E, de Rossi P, Bacioglu M, Langston RG, Kaganovich A, Bengoa-Vergniory N, Gonzalez-Guerra A, Avar M, Heinzer D, Reimann R, Hasler LM, Herling TW, Matharu NS, Landeck N, Luk K, Melki R, Kahle PJ, Hornemann S, Knowles TPJ, Cookson MR, Polymenidou M, Jucker M, Aguzzi A. LAG3 is not expressed in human and murine neurons and does not modulate alpha-synucleinopathies. EMBO Mol Med. 2021;13(9):e14745. Epub 20210726. doi: 10.15252/emmm.202114745. PubMed PMID: 34309222; PMCID: PMC8422075.

15. Carithers LJ, Ardlie K, Barcus M, Branton PA, Britton A, Buia SA, Compton CC, DeLuca DS, Peter-Demchok J, Gelfand ET, Guan P, Korzeniewski GE, Lockhart NC, Rabiner CA, Rao AK, Robinson KL, Roche NV, Sawyer SJ, Segre AV, Shive CE, Smith AM, Sobin LH, Undale AH, Valentino KM, Vaught J, Young TR, Moore HM, Consortium GT. A Novel Approach to High-Quality Postmortem Tissue Procurement: The GTEx Project. Biopreserv Biobank. 2015;13(5):311–9. doi: 10.1089/bio.2015.0032. PubMed PMID: 26484571; PMCID: PMC4675181.

16. Consortium GT. The Genotype-Tissue Expression (GTEx) project. Nat Genet. 2013;45(6):580–5. doi: 10.1038/ng.2653. PubMed PMID: 23715323; PMCID: PMC4010069.

17. Lein ES, Hawrylycz MJ, Ao N, Ayres M, Bensinger A, Bernard A, Boe AF, Boguski MS, Brockway KS, Byrnes EJ, Chen L, Chen L, Chen TM, Chin MC, Chong J, Crook BE, Czaplinska A, Dang CN, Datta S, Dee NR, Desaki AL, Desta T, Diep E, Dolbeare TA, Donelan MJ, Dong HW, Dougherty JG, Duncan BJ, Ebbert AJ, Eichele G, Estin LK, Faber C, Facer BA, Fields R, Fischer SR, Fliss TP, Frensley C, Gates SN, Glattfelder KJ, Halverson KR, Hart MR, Hohmann JG, Howell MP, Jeung DP, Johnson RA, Karr PT, Kawal R, Kidney JM, Knapik RH, Kuan CL, Lake JH, Laramee AR, Larsen KD, Lau C, Lemon TA, Liang AJ, Liu Y, Luong LT, Michaels J, Morgan JJ, Morgan RJ, Mortrud MT, Mosqueda NF, Ng LL, Ng R, Orta GJ, Overly CC, Pak TH, Parry SE, Pathak SD, Pearson OC, Puchalski RB, Riley ZL, Rockett HR, Rowland SA, Royall JJ, Ruiz MJ, Sarno NR, Schaffnit K, Shapovalova NV, Sivisay T, Slaughterbeck CR, Smith SC, Smith KA, Smith BI, Sodt AJ, Stewart NN, Stumpf KR, Sunkin SM, Sutram M, Tam A, Teemer CD, Thaller C, Thompson CL, Varnam LR, Visel A, Whitlock RM, Wohnoutka PE, Wolkey CK, Wong VY, Wood M, Yaylaoglu MB, Young RC, Youngstrom BL, Yuan XF, Zhang B, Zwingman TA, Jones AR. Genome-wide atlas of gene expression in the adult mouse brain. Nature. 2007;445(7124):168-76. Epub 20061206. doi: 10.1038/nature05453. PubMed PMID: 17151600.

18. Moreno P, Fexova S, George N, Manning JR, Miao Z, Mohammed S, Munoz-Pomer A, Fullgrabe A, Bi Y, Bush N, Iqbal H, Kumbham U, Solovyev A, Zhao L, Prakash A, Garcia-Seisdedos D, Kundu DJ, Wang S, Walzer M, Clarke L, Osumi-Sutherland D, Tello-Ruiz MK, Kumari S, Ware D, Eliasova J, Arends MJ, Nawijn MC, Meyer K, Burdett T, Marioni J, Teichmann S, Vizcaino JA, Brazma A, Papatheodorou I. Expression Atlas update: gene and protein expression in multiple species. Nucleic Acids Res. 2022;50(D1):D129–D40. doi: 10.1093/nar/gkab1030. PubMed PMID: 34850121; PMCID: PMC8728300.

19. Papatheodorou I, Moreno P, Manning J, Fuentes AM, George N, Fexova S, Fonseca NA, Fullgrabe A, Green M, Huang N, Huerta L, Iqbal H, Jianu M, Mohammed S, Zhao L, Jarnuczak AF, Jupp S, Marioni J, Meyer K, Petryszak R, Prada Medina CA, Talavera-Lopez C, Teichmann S, Vizcaino JA, Brazma A. Expression Atlas update: from tissues to single cells. Nucleic Acids Res. 2020;48(D1):D77–D83. doi: 10.1093/nar/gkz947. PubMed PMID: 31665515; PMCID: PMC7145605.

20. Wu C, Jin X, Tsueng G, Afrasiabi C, Su AI. BioGPS: building your own mash-up of gene annotations and expression profiles. Nucleic Acids Res. 2016;44(D1):D313–6. Epub 20151117. doi: 10.1093/nar/gkv1104. PubMed PMID: 26578587; PMCID: PMC4702805.

21. Wu C, Orozco C, Boyer J, Leglise M, Goodale J, Batalov S, Hodge CL, Haase J, Janes J, Huss JW, 3rd, Su AI. BioGPS: an extensible and customizable portal for querying and organizing gene annotation resources. Genome Biol. 2009;10(11):R130. Epub 20091117. doi: 10.1186/gb-2009-10-11-r130. PubMed PMID: 19919682; PMCID: PMC3091323.

22. Chen C, Kumbhar R, Wang H, Yang X, Gadhave K, Rastegar C, Kimura Y, Behensky A, Kotha S, Kuo G, Katakam S, Jeong D, Wang L, Wang A, Chen R, Zhang S, Jin L, Workman CJ, Vignali DAA, Pletinkova O, Jia H, Peng W, Nauen DW, Wong PC, Redding-Ochoa J, Troncoso JC, Ying M, Dawson VL, Dawson TM, Mao X. Lymphocyte-Activation Gene 3 Facilitates Pathological Tau Neuron-to-Neuron Transmission. Adv Sci (Weinh). 2024;11(16):e2303775. Epub 20240207. doi: 10.1002/advs.202303775. PubMed PMID: 38327094; PMCID: PMC11040377.

23. Zhang Q, Duan Q, Gao Y, He P, Huang R, Huang H, Li Y, Ma G, Zhang Y, Nie K, Wang L. Cerebral Microvascular Injury Induced by Lag3-Dependent alpha-Synuclein Fibril Endocytosis Exacerbates Cognitive Impairment in a Mouse Model of alpha-Synucleinopathies. Adv Sci (Weinh). 2023;10(25):e2301903. Epub 20230628. doi: 10.1002/advs.202301903. PubMed PMID: 37381656; PMCID: PMC10477873.

24. Zhang Q, Chikina M, Szymczak-Workman AL, Horne W, Kolls JK, Vignali KM, Normolle D, Bettini M, Workman CJ, Vignali DAA. LAG3 limits regulatory T cell proliferation and function in autoimmune diabetes. Sci Immunol. 2017;2(9). doi: 10.1126/sciimmunol.aah4569. PubMed PMID: 28783703; PMCID: PMC5609824.

25. Tronche F, Kellendonk C, Kretz O, Gass P, Anlag K, Orban PC, Bock R, Klein R, Schutz G. Disruption of the glucocorticoid receptor gene in the nervous system results in reduced anxiety. Nat Genet. 1999;23(1):99–103. doi: 10.1038/12703. PubMed PMID: 10471508.

26. Giusti SA, Vercelli CA, Vogl AM, Kolarz AW, Pino NS, Deussing JM, Refojo D. Behavioral phenotyping of Nestin-Cre mice: implications for genetic mouse models of psychiatric disorders. J Psychiatr Res. 2014;55:87–95. Epub 20140412. doi: 10.1016/j.jpsychires.2014.04.002. PubMed PMID: 24768109.

27. Volpicelli-Daley LA, Luk KC, Lee VM. Addition of exogenous alpha-synuclein preformed fibrils to primary neuronal cultures to seed recruitment of endogenous alpha-synuclein to Lewy body and Lewy neurite-like aggregates. Nat Protoc. 2014;9(9):2135–46. Epub 20140814. doi: 10.1038/nprot.2014.143. PubMed PMID: 25122523; PMCID: PMC4372899.

28. Thi Lai T, Kim YE, Nguyen LTN, Thi Nguyen T, Kwak IH, Richter F, Kim YJ, Ma HI. Microglial inhibition alleviates alpha-synuclein propagation and neurodegeneration in Parkinson’s disease mouse model. NPJ Parkinsons Dis. 2024;10(1):32. Epub 20240202. doi: 10.1038/s41531-024-00640-2. PubMed PMID: 38302446; PMCID: PMC10834509.

29. Kim S, Kwon SH, Kam TI, Panicker N, Karuppagounder SS, Lee S, Lee JH, Kim WR, Kook M, Foss CA, Shen C, Lee H, Kulkarni S, Pasricha PJ, Lee G, Pomper MG, Dawson VL, Dawson TM, Ko HS. Transneuronal Propagation of Pathologic alpha-Synuclein from the Gut to the Brain Models Parkinson’s Disease. Neuron. 2019;103(4):627–41 e7. Epub 20190626. doi: 10.1016/j.neuron.2019.05.035. PubMed PMID: 31255487; PMCID: PMC6706297.

30. Eninger T, Muller SA, Bacioglu M, Schweighauser M, Lambert M, Maia LF, Neher JJ, Hornfeck SM, Obermuller U, Kleinberger G, Haass C, Kahle PJ, Staufenbiel M, Ping L, Duong DM, Levey AI, Seyfried NT, Lichtenthaler SF, Jucker M, Kaeser SA. Signatures of glial activity can be detected in the CSF proteome. Proc Natl Acad Sci U S A. 2022;119(24):e2119804119. Epub 20220606. doi: 10.1073/pnas.2119804119. PubMed PMID: 35666874; PMCID: PMC9214531.

31. Tozzi A, de Iure A, Bagetta V, Tantucci M, Durante V, Quiroga-Varela A, Costa C, Di Filippo M, Ghiglieri V, Latagliata EC, Wegrzynowicz M, Decressac M, Giampa C, Dalley JW, Xia J, Gardoni F, Mellone M, El-Agnaf OM, Ardah MT, Puglisi-Allegra S, Bjorklund A, Spillantini MG, Picconi B, Calabresi P. Alpha-Synuclein Produces Early Behavioral Alterations via Striatal Cholinergic Synaptic Dysfunction by Interacting With GluN2D N-Methyl-D-Aspartate Receptor Subunit. Biol Psychiatry. 2016;79(5):402–14. Epub 20150820. doi: 10.1016/j.biopsych.2015.08.013. PubMed PMID: 26392130.

32. Diogenes MJ, Dias RB, Rombo DM, Vicente Miranda H, Maiolino F, Guerreiro P, Nasstrom T, Franquelim HG, Oliveira LM, Castanho MA, Lannfelt L, Bergstrom J, Ingelsson M, Quintas A, Sebastiao AM, Lopes LV, Outeiro TF. Extracellular alpha-synuclein oligomers modulate synaptic transmission and impair LTP via NMDA-receptor activation. J Neurosci. 2012;32(34):11750–62. doi: 10.1523/JNEUROSCI.0234-12.2012. PubMed PMID: 22915117; PMCID: PMC6703775.

33. Li S, Jin M, Koeglsperger T, Shepardson NE, Shankar GM, Selkoe DJ. Soluble Abeta oligomers inhibit long-term potentiation through a mechanism involving excessive activation of extrasynaptic NR2B-containing NMDA receptors. J Neurosci. 2011;31(18):6627–38. doi: 10.1523/JNEUROSCI.0203-11.2011. PubMed PMID: 21543591; PMCID: PMC3100898.

34. Ferreira DG, Temido-Ferreira M, Vicente Miranda H, Batalha VL, Coelho JE, Szego EM, Marques-Morgado I, Vaz SH, Rhee JS, Schmitz M, Zerr I, Lopes LV, Outeiro TF. alpha-synuclein interacts with PrP(C) to induce cognitive impairment through mGluR5 and NMDAR2B. Nat Neurosci. 2017;20(11):1569–79. Epub 20170925. doi: 10.1038/nn.4648. PubMed PMID: 28945221.

35. Sagal J, Zhan X, Xu J, Tilghman J, Karuppagounder SS, Chen L, Dawson VL, Dawson TM, Laterra J, Ying M. Proneural transcription factor Atoh1 drives highly efficient differentiation of human pluripotent stem cells into dopaminergic neurons. Stem Cells Transl Med. 2014;3(8):888–98. Epub 20140605. doi: 10.5966/sctm.2013-0213. PubMed PMID: 24904172; PMCID: PMC4116248.

36. Cheng HJ, Flanagan JG. Identification and cloning of ELF-1, a developmentally expressed ligand for the Mek4 and Sek receptor tyrosine kinases. Cell. 1994;79(1):157–68. doi: 10.1016/0092-8674(94)90408-1. PubMed PMID: 7522971.

37. Cheng HJ, Flanagan JG. Cloning and characterization of RTK ligands using receptor- alkaline phosphatase fusion proteins. Methods Mol Biol. 2001;124:313–34. doi: 10.1385/1-59259-059-4:313. PubMed PMID: 11100484.

38. Workman CJ, Cauley LS, Kim IJ, Blackman MA, Woodland DL, Vignali DA. Lymphocyte activation gene-3 (CD223) regulates the size of the expanding T cell population following antigen activation in vivo. J Immunol. 2004;172(9):5450–5. doi: 10.4049/jimmunol.172.9.5450. PubMed PMID: 15100286.

39. West MJ. New stereological methods for counting neurons. Neurobiol Aging. 1993;14(4):275–85. doi: 10.1016/0197-4580(93)90112-o. PubMed PMID: 8367009.

40. Lee Y, Karuppagounder SS, Shin JH, Lee YI, Ko HS, Swing D, Jiang H, Kang SU, Lee BD, Kang HC, Kim D, Tessarollo L, Dawson VL, Dawson TM. Parthanatos mediates AIMP2- activated age-dependent dopaminergic neuronal loss. Nat Neurosci. 2013;16(10):1392–400. Epub 20130825. doi: 10.1038/nn.3500. PubMed PMID: 23974709; PMCID: PMC3785563.

41. Ting JT, Daigle TL, Chen Q, Feng G. Acute brain slice methods for adult and aging animals: application of targeted patch clamp analysis and optogenetics. Methods Mol Biol. 2014;1183:221–42. doi: 10.1007/978-1-4939-1096-0_14. PubMed PMID: 25023312; PMCID: PMC4219416.

42. Bywood PT, Johnson SM. Differential vulnerabilities of substantia nigra catecholamine neurons to excitatory amino acid-induced degeneration in rat midbrain slices. Exp Neurol. 2000;162(1):180–8. doi: 10.1006/exnr.2000.7310. PubMed PMID: 10716898.

43. VanderHorst VG, Ulfhake B. The organization of the brainstem and spinal cord of the mouse: relationships between monoaminergic, cholinergic, and spinal projection systems. J Chem Neuroanat. 2006;31(1):2–36. Epub 20050923. doi: 10.1016/j.jchemneu.2005.08.003. PubMed PMID: 16183250.

44. Wanat MJ, Bonci A. Dose-dependent changes in the synaptic strength on dopamine neurons and locomotor activity after cocaine exposure. Synapse. 2008;62(10):790–5. doi: 10.1002/syn.20546. PubMed PMID: 18655120; PMCID: PMC5018993.

45. Zhang TA, Placzek AN, Dani JA. In vitro identification and electrophysiological characterization of dopamine neurons in the ventral tegmental area. Neuropharmacology. 2010;59(6):431–6. Epub 20100618. doi: 10.1016/j.neuropharm.2010.06.004. PubMed PMID: 20600174; PMCID: PMC2946471.

46. Amendola J, Woodhouse A, Martin-Eauclaire MF, Goaillard JM. Ca(2)(+)/cAMP-sensitive covariation of I(A) and I(H) voltage dependences tunes rebound firing in dopaminergic neurons. J Neurosci. 2012;32(6):2166–81. doi: 10.1523/JNEUROSCI.5297-11.2012. PubMed PMID: 22323729; PMCID: PMC6621702.

